# Confluence of timing and reward biases in perceptual decision-making dynamics

**DOI:** 10.1101/865501

**Authors:** Maxwell Shinn, Daniel Ehrlich, Daeyeol Lee, John D. Murray, Hyojung Seo

**Affiliations:** Department of Psychiatry, New Haven, CT, USA; Department of Neuroscience, Yale University, New Haven, CT, USA; Interdepartmental Neuroscience Program, Yale University, New Haven, CT, USA

## Abstract

Although the decisions of our daily lives often occur in the context of temporal and reward structures, the impact of such regularities on decision-making strategy is poorly understood. Here, to explore how temporal and reward context modulate strategy, we trained rhesus monkeys to perform a novel perceptual decision-making task with asymmetric rewards and time-varying evidence reliability. To model the choice and response time patterns, we developed a computational framework for fitting generalized drift-diffusion models (GDDMs) which flexibly accommodates diverse evidence accumulation strategies. We found that a dynamic urgency signal and leaky integration, in combination with two independent forms of reward biases, best capture behavior. We also tested how temporal structure influences urgency by systematically manipulating the temporal structure of sensory evidence, and found that the time course of urgency was affected by temporal context. Overall, our approach identified key components of cognitive mechanisms for incorporating temporal and reward structure into decisions.

## Introduction

In an uncertain and dynamic environment, humans and other animals detect temporal regularities in the environment and use them to inform decision-making (***Woodrow, 1914; Rosenbaum and Collyer, 1998; Los, 2010; Behrens et al., 2007; Farashahi et al., 2017***). Even in relatively simple perceptual tasks, the timing and accuracy of decisions are sensitive to the temporal statistics of stimuli (***Grosjean et al., 2001***). Cognitive strategies in perceptual tasks are often reflected in the response times (RTs), which therefore can be effectively used to test mechanistic models of decision-making processes (***Luce, 1986***).

One paradigm for studying perceptual decision-making computations and their neural correlates is to present dynamic sensory evidence over time (***Newsome et al., 1989; Roitman and Shadlen, 2002***). Evidence accumulation has been proposed as a leading strategy for decision-making under this paradigm, which can be formalized using the drift-diffusion model (DDM) (***Ratcliff, 1978; Ratcliff et al., 2016***). The DDM has been employed to capture choice and RT behavior in a range of decision-making tasks (***Palmer et al., 2005; Krajbich and Rangel, 2011; Ding and Gold, 2012; Bogacz et al., 2006; Resulaj et al., 2009; Ratcliff et al., 2003; Mormann et al., 2010***). In the DDM, a dynamic decision variable integrates evidence over time, and a decision is reached when this variable crosses a bound. An open question is how evidence accumulation is shaped by temporal uncertainty or expectation of the stimulus. For instance, if the signal-to-noise ratio changes across the time of stimulus presentation, can the integration process dynamically gate this input signal?

Such gating is important for integrating relevant information while ignoring irrelevant information. Uncertainty in the temporal onset of information creates a need to modulate the integration process over time. For example, beginning integration before relevant information is available causes noise to be unnecessarily integrated, thereby reducing decision accuracy (***Laming, 1979; Grosjean et al., 2001***). By contrast, beginning integration after the onset of the stimulus discards potentially beneficial information (***Devine et al., 2019***). It is unknown how these timing factors should be implemented within a DDM framework. Time-varying bounds (***Murphy et al., 2016; Hanks et al., 2014; Ditterich, 2006b; Drugowitsch et al., 2012***) and gain modulation (***Ditterich, 2006b; Cisek et al., 2009; Thura et al., 2012***) have improved predictive ability for RT patterns, but these mechanisms have not been utilized to explain uncertainty in stimulus onset time. Furthermore, the utility of DDMs incorporating these and other complex extensions is limited by the computational challenges of simulating and fitting such models to empirical choice and RT patterns.

In addition to sensory evidence and temporal expectation, decisions should incorporate differences in value among the options. Reward bias has long been an active topic of research (***Feng et al., 2009; Rorie et al., 2010; Laming, 1968; Ratcliff, 1985; Mulder et al., 2012***), with more recent attention to its connections to timing (***Drugowitsch et al., 2012; Voskuilen et al., 2016; Hanks et al., 2011; Lauwereyns et al., 2002; Nagano-Saito et al., 2012; Gao et al., 2011***). While these asymmetries have previously been incorporated into the DDM framework (***Ratcliff and McKoon, 2008; Ratcliff, 1985; Edwards, 1965; Laming, 1968; Mulder et al., 2012***), the strategies through which sensory, value, and timing information are combined to form a decision are still poorly understood.

To investigate these issues, we developed a novel behavioral paradigm for perceptual decision-making in which the onset of the evidence is temporally uncertain. Therefore, the temporal structure of evidence could be strategically exploited during decision-making. By simultaneously manipulating reward differences between options, we also examined whether and how time-varying evidence interacts with temporal strategies to produce reward bias. We trained monkeys to perform this task, and found that the animals exploited the temporal structure of the evidence and adjusted the timing of their decisions in such a way to reduce uncertainty about the evidence onset.

To quantitatively model the decision-making behavior, we developed a framework for computationally efficient simulation and fitting of generalized DDMs (GDDMs) to choice and RT behavior. GDDMs encompass both the DDM and also non-integrative models as special cases (***Cisek et al., 2009; Thura et al., 2012***). We found that a GDDM with timing and reward processes can quantitatively capture the animal’s behavior as determined by its RT distribution. Temporal modulation of the integration process can be implemented through an urgency signal which varies across time. Such an urgency signal can flexibly adjust the animal’s behavior as task timing demands change. In addition, capturing reward-related behavioral effects required two mechanisms, one dependent on the integration process and the other independent of integration. Overall, our findings suggest that evidence accumulation in perceptual decision-making can be flexibly modulated by temporal and reward expectation, and that generalized DDMs can mathematically capture these phenomena.

## Results

### Behavioral task

Two rhesus monkeys (*Macaca mulatta*) were trained to perform a two-alternative forced choice, color matching task (***Figure 1***). In each trial, a central square patch was presented consisting of a 20×20 grid of green and blue pixels that rearranged randomly at 20 Hz. Stimulus presentation was divided into two consecutive periods containing an uninformative “presample” and informative “sample”. The animal indicated its choice by shifting its gaze to one of two flanking choice targets, one green and one blue. The trial was rewarded via juice delivery if the selected target color corresponded to the majority color of pixels in the sample. Task difficulty was manipulated by parametrically varying the proportion of pixels of each color in the sample, which we refer to as color coherence. Animals were allowed to direct their gaze to a choice target any time after the onset of the sample. In addition, reward cues were displayed surrounding the saccade targets which indicated whether a large or small reward would be delivered for a correct response to the corresponding target. These reward cues were randomly assigned to a target on each trial. Zero coherence corresponds to a 50/50 mixture of blue and green pixels, and coherence of ±1 corresponds to a solid color. By convention, we define coherence relative to large- and small-reward targets, where a coherence of+ 1 or −1 corresponds to all pixels being the color of the large-reward or small-reward target, respectively.

**Figure 1.**
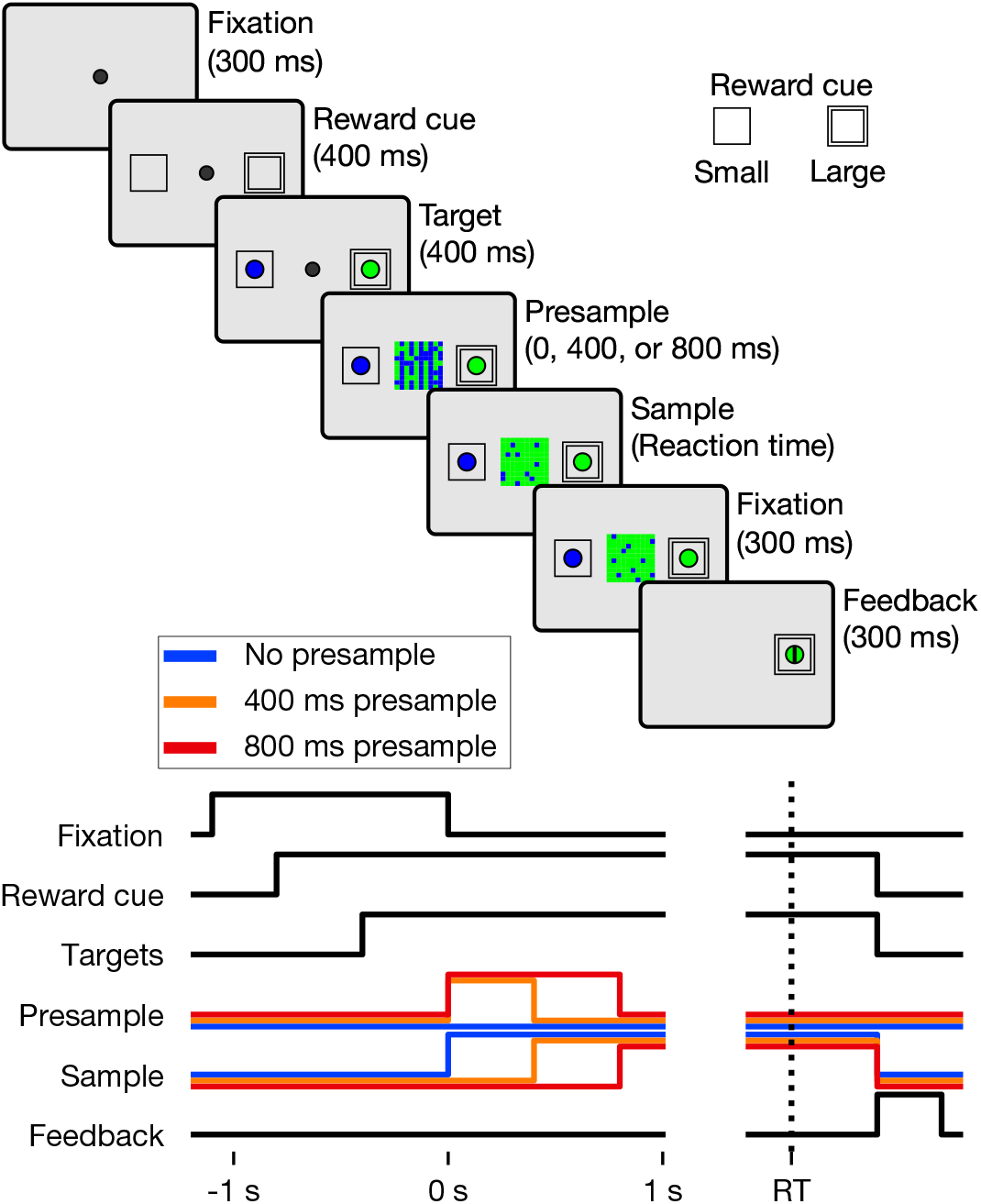
Color matching task with asymmetric reward. The temporal sequence of trial events in the color matching task with asymmetric reward is shown above. Reward cues indicating a large or small reward are inset. Below is a timeline indicating the presence of various task elements on the screen, with presample and sample lines for each condition denoted by the line color. **Figure 1–video 1. Video of the task.** The task is demonstrated across several coherences and presamples. Element sizes are adjusted for demonstration.

Temporal uncertainty in the sensory evidence was introduced by varying the duration of the uninformative presample. No explicit cue was presented to indicate this transition from presample to sample. During the presample period, the color coherence was zero, and at the transition to the sample, the coherence switched to the particular value chosen for that trial. The presample duration was selected randomly from three possible time intervals—0, 0.4 or 0.8 s—with equal probability. A premature choice was punished by a 2 s timeout. In a separate experiment described below, we manipulated the set of presample durations (***Figure 6***).

### Task parameters influence behavior

We predicted that all three primary task variables—coherence, presample duration, and reward location—would lead to systematic changes in the animal’s behavior as measured by the RT distributions. In this study, RT was measured from the onset of the presample, since the transition from presample to sample was not explicitly cued. To test the impact of these three task variables on the animal’s performance, we used multiple-regression models to predict RT and accuracy in individual trials using these three task variables (see Methods).

We found that all three task variables had a significant influence on RT (*p* < .05 for both animals, ***Equation 1***) and accuracy (*p* < .05 for both animals, ***Equation 3***). Higher absolute coherences decreased the RT and increased accuracy, meaning that animals responded faster (***Figure 2***b,e) and more accurately (***Figure 2***a,d) on easier trials. Likewise, responses directed towards the large-reward target were faster than those directed at the small-reward target (***Figure 2***b,e). Furthermore, responses were faster and more accurate on trials with shorter presample durations (***Figure 2***a,b,d,e).

**Figure 2.**
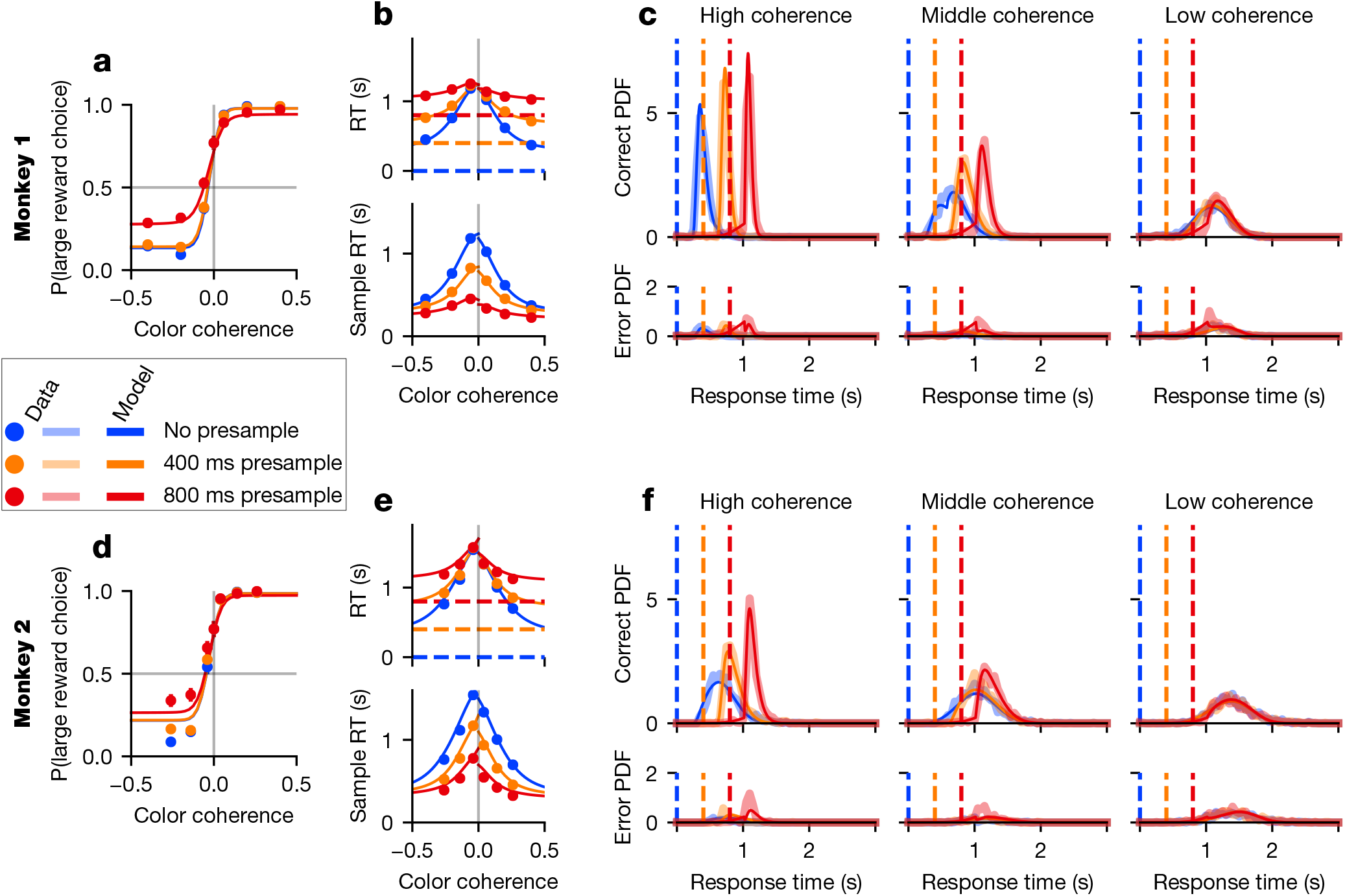
Animal and model performance. Psychometric (a,d) and chronometric (b,e) functions, as well as RT histograms (c,f), are plotted for the validation data of Monkey 1 (a,b,c) and 2 (d,e,f). Data are plotted for each coherence value as dots in the behavioral function (a,b,d,e) and as thick translucent lines in the probability distributions (c,f). “Sample RT” indicates the RT minus the presample duration. Overlaid as a thin solid line is the best-fit generalized drift-diffusion model described in section GDDM accounts for strategies with parameters fit to the exploration data. Error bars for 95% CI are hidden beneath the markers. **Figure 2–Figure supplement 1. DDM with reward bias does not capture monkey behavior.** Format similar to ***Figure 2***, but with the model described in Drift-diffusion model with reward mechanisms. **Figure 2–Figure supplement 2. Animal and model performance in exploration dataset.** Format similar to ***Figure 2***, but showing the exploration data using model parameters fit to it.

We next examined whether these task variables modulated RT and accuracy independently. We found that both accuracy (*p* < .05 for both animals, ***Equation 2***) and RT (*p* < .05 for both animals, ***Equation 4***) were significantly modulated by the interaction between absolute coherence and presample duration. This interaction’s effect on RT can be seen in the chronometric function (***Figure 2***b,e) by observing that for low-coherence trials, RT aligned to the presample is approximately equal regardless of the presample duration. In other words, despite the fact that there is 800ms of additional sensory evidence presented to the animal during the 0ms presample trials compared to the 800ms presample trials, the RT is similar in both cases. Such similarity in response time is reflected in the overlap in the RT distribution in low-coherence trials for all presample durations (***Figure 2***c,f). By contrast, for middle- or high-coherence trials, the RT shows a larger effect of coherence.

In summary, we find that coherence, presample duration, and large-reward location not only modulated RT and accuracy independently, but the interaction between coherence and presample duration influences choice and RT. To gain insights into the underlying mechanisms, we tested whether a DDM that incorporated these task variables could provide a parsimonious account for the observed behaviors.

### Drift-diffusion model with reward mechanisms

The DDM is one of the simplest models that formalizes evidence integration for two alternative choices (***Luce, 1986; Bogacz et al., 2006; Ratcliff et al., 2016***). Recently, studies have shown that reward bias in psychometric functions can be accounted for by two separate mechanisms (***Fan et al., 2018; Gesiarz et al., 2019***). First, the initial bias shifts the starting position of the decision variable in the direction of the large-reward target (***Edwards, 1965; Laming, 1968; Ratcliff, 1985***). Second, a time-dependent bias allows a constant evidence signal to be continuously added to the integrator in the direction of the large-reward choice (***Mulder et al., 2012; Ratcliff, 1985; Ashby, 1983***). These two types of reward biases have been shown to be effective in reproducing choice bias (***van Ravenzwaaij et al., 2012; Voss et al., 2004; Diederich and Busemeyer, 2006; Ratcliff and McKoon, 2008***).

We thus took this combined reward-biased DDM (***Fan et al., 2018; Gesiarz et al., 2019***) as a point of departure to gain further insights and understanding of our data. In this extended seven-parameter model (see Methods, ***Equation 7***), the weight on evidence is constant over time, as are the bounds. We fit this model by maximizing likelihood using the method of differential evolution (***Storn and Price, 1997***).

While the DDM with combined reward bias correctly predicted the large-reward target would be chosen with a higher probability and a shorter RT than the small-reward target, overall this model could not fit other major features of the choice and RT data as measured by the psychometric (***Figure 2–Figure Supplement 1***a,d) and chronometric functions (***Figure 2–Figure Supplement 1***b,e) as well as the RT distributions (***Figure 2–Figure Supplement 1***c,f). First, it failed to reproduce the overlapping RT distributions across different durations of presample for low-coherence samples (***Figure 2–Figure Supplement 1***c,f). This model, like DDM models in general, predicts that RT should depend on the amount of evidence which has been integrated. However, the overlapping RT distributions indicate a minimal effect of evidence integrated during the early part of the trial compared to evidence later in the trial.

Second, the model failed to produce the characteristic shapes of RT distributions in the data. In both the data and the model, color coherence and sample time jointly influenced the shape of the RT distribution. However, the distribution of the actual RT data became more symmetric as color coherence decreased, whereas the distribution of RT in the model remained skewed similarly in all conditions (***Figure 2–Figure Supplement 1***c,f top panels).

Finally, the model failed to fit the time course of errors induced by the strong reward bias. During the task, errors during high-coherence stimuli were made almost exclusively because animals tended to choose the large-reward target when the evidence supported the small-reward choice (***Figure 2–Figure Supplement 1***a,d). Therefore, errors during the high-coherence sample particularly reflect animals’ strong bias toward large-reward target and the error rate tended to sharply increase immediately after the sample onset (***Figure 2–Figure Supplement 1***c,f bottom panels). By contrast, the DDM predicted that fewer errors must follow the onset of the sample, as more evidence supporting correct choice becomes available after sample onset. These results collectively suggest that animals’ behavior during the task used in our study cannot be fully explained by a time-invariant evidence integration strategy in conjunction with simple extensions to incorporate reward bias.

### Elements of Generalized DDMs

The above results suggest that relatively simple modifications of the DDM are insufficient in that they cannot accurately reproduce the RT distribution. This is especially true of their ability to model the evolution of reward effects over time under temporally uncertain conditions. Therefore, we extended the DDM framework to create a generalized drift-diffusion model (GDDM) for testing potential mechanisms to account for animals’ behavior in our task. GDDMs allow the drift rate and diffusion coefficient to be arbitrary functions of time and of the position of integrated evidence (i.e. decision variable), and integration bounds to be arbitrary functions of time. GDDMs also allow for a leaky integrator, meaning the model can incorporate both integrative and non-integrative strategies that utilize instantaneous evidence (***Cisek et al., 2009; Thura et al., 2012***) (see Methods, ***Equation 9*** for the details of the model). Therefore, the GDDM enables fitting of drift-diffusion models that implement complex time-varying mechanisms for evidence sampling and integration as well as reward bias.

#### Mechanisms for modulating the time scale of evidence integration

While the simple DDM assumes perfect integration in which the integrated evidence does not decay over time (i.e. “forgetting”), we consider the possibility that animals might integrate evidence at a relatively short time scale so that the integrated evidence primarily reflects the most recent evidence (***Usher and McClelland, 2001; Cisek et al., 2009; Brown and Holmes, 2001; Feng et al., 2009; Brown et al., 2005; Ossmy et al., 2013; Veliz-Cuba et al., 2016***). Therefore, the GDDM included a “leak” parameter that determines the time constant of the decay in decision variable. A leak parameter permits the model to capture both perfect integration, where all evidence is considered in the decision-making process, and non-integration, where only momentary evidence is considered (***Cisek et al., 2009***), thereby generalizing across these distinct strategies.

#### Mechanisms of time-varying urgency

To explain animals’ strategic timing of decisions, we hypothesized that animals might dynamically modulate the evidence accumulation process as a function of time during the trial. Within the DDM framework, this was implemented through a time dependence of parameter values, which has been most studied in the context of an “urgency” signal. An urgency signal can be characterized across two dimensions: its type, and its form. Two types of urgency signal include decision bounds which collapse over time (***Ditterich, 2006b; Drugowitsch et al., 2012; Hanks et al., 2014; Murphy et al., 2016***) and a time-dependent increase in the gain of the momentary sensory evidence (***Cisek et al., 2009; Gold and Shadlen, 2001; Ditterich, 2006b***). Here, we consider both of these time-varying urgency signal types as means to modulate the evidence accumulation process.

In addition to the urgency signal’s type, we can also consider its form. Consistent with prior work, animals might have begun increasing their urgency at the beginning of the trial. We implemented this as a linearly increasing gain function (“linear gain”, ***Figure 3***a) or a gradually decreasing bound (“collapsing bounds”, ***Figure 3***a). The limiting cases—where the slope of the gain function is zero or the time constant of the collapsing bound is infinite—correspond to time-independent urgency, or equivalently to the lack of explicit urgency in the simple DDM (“constant”, ***Figure 3***a).

**Figure 3.**
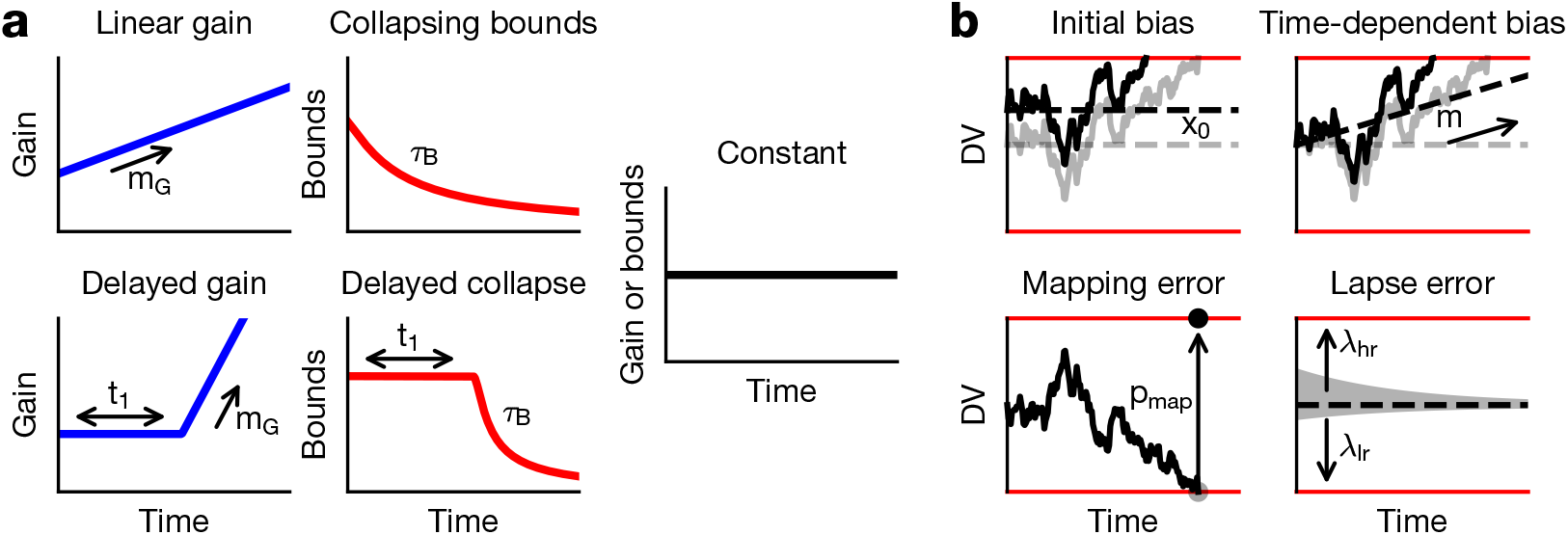
Generalized DDM components. Our GDDM included extensions related to timing and reward. (a) We modeled timing using two types of urgency signals. We implemented a gain function and collapsing bounds, each with and without a time delay. Constant gain and constant bounds indicates the absence of an urgency signal. Bounds are shown in red, and gain functions in blue. (b) Reward mechanisms are shown with example decision variable trajectories for each mechanism. Initial bias: the integrator starts biased towards the large-reward target, and the leaky integrator decays back to this starting position instead of to the origin. Time-dependent bias: there is a gradual increase in baseline evidence towards the large-reward target over time. Mapping error: once a decision is reached, the monkey chooses the opposite target on a percentage of trials. Lapse error: there is a higher, exponentially-distributed probability of making an evidence-independent choice to the large-reward target at any given point throughout the trial, contrasted to equal probabilities in the absence of this mechanism.

In a simple form of a task-specific time-varying urgency signal, animals might have delayed increasing urgency until the uncertainty of evidence onset was sufficiently resolved. Therefore, in addition to urgency signals that begin to ramp at the onset of the sensory stimulus, as is commonly described in the literature, we additionally considered urgency signals which do not begin to ramp until some later point within the trial. We tested both gain modulation and collapsing bounds urgency signals with such a nonlinear time course to allow for an effect of task structure on urgency. The delayed urgency signal introduced a delay before the linear increase in the gain function (“delayed gain”, ***Figure 3***a) or exponential collapse of the bounds (“delayed collapse”, ***Figure 3***a). This delay parameter would be tuned by the temporal uncertainty of the behavioral task.

#### Mechanisms of reward bias

We found that animals’ bias toward the large-reward choice increased as the sample was presented at more predictable time, particularly for higher coherence samples (***Figure 2***a,d). We considered two classes of mechanism, one in which reward directly influences the integration process, and the other in which the reward bias is implemented outside the integration process.

Similar to how the DDM was extended to incorporate reward bias, the effect of reward on the integration process can be modeled by an “initial bias”, which maintains a fixed magnitude throughout the trial, or a “time-dependent bias”, which increases in magnitude throughout the trial (***Fan et al., 2018; Gesiarz et al., 2019***) (***Figure 2–Figure Supplement 1***; ***Figure 3***b). The initial bias is traditionally implemented as an initial value of the decision variable, but due to leaky integration, this is ineffective because the decision variable will decay from this starting position back to zero. Thus, in addition to setting the initial value of the decision variable, our implementation of initial bias also causes the decision variable to leak towards this initial position. For similar reasons, our implementation of the time-dependent bias is a linear increase in the value to which the decision variable leaks over the course of the trial (see Methods).

We also considered two additional types of reward bias which are outside the integration process. First, the animal may be biased in its categorical choice at the end of evidence accumulation (***Erlich et al., 2015; Hanks et al., 2015***). Namely, when the decision variable reaches the bound, animals may with some probability mistakenly produce a motor response towards the large-reward choice even if they correctly reached the bound for small-reward choice (“mapping error”, ***Figure 3***b). Second, we also considered that a small number of response may be generated randomly anytime during the integration process (“lapse trials”). The responses on these lapse trials may be biased toward the large-reward choice (***Simen et al., 2009; Noorbaloochi et al., 2015***), which can be implemented by an asymmetric lapse rate for the large- and small-reward choices (“lapse bias”, ***Figure 3***b) (***Yartsev et al., 2018***).

#### Robust estimation of model parameters

As the models described here have many parameters, methods for parameter estimation must be carefully considered. First, we must consider the metric by which potential models are evaluated. We used a state-of-the-art simulation environment which allows fitting models using maximum likelihood on the full probability distribution. Further details are described in Section Fitting method.

Second, we must protect against overfitting. Before performing any analyses on the data for the task with asymmetric reward, the data were split in half by pseudo-randomly choosing half of the trials for the “validation” set and excluding these from further analysis, analyzing only the disjoint “exploration” data. The validation trials were not fit to the model or otherwise examined until all analyses for the present manuscript were complete. After unmasking the validation trials, no additional analyses were performed on the data. More details are described in Section Cross-validation procedure.

### GDDM accounts for strategies

We fit the GDDM to the choice and RT behavior for each animal, and evaluated model fit with held-out log-likelihood (HOLL). Overall, models with time-varying urgency performed remarkably better than ones with constant urgency such as the simple DDM (***Figure 4***a,b). When combined with each of several different reward bias mechanisms, models with delayed urgency best explained the data for monkey 1, while models with non-delayed and delayed urgency performed similarly well for monkey 2. Despite some individual differences, these results suggest that the animal’s strategic adaptation to the temporal structure of sensory evidence can be well accounted for by a time-varying urgency.

**Figure 4.**
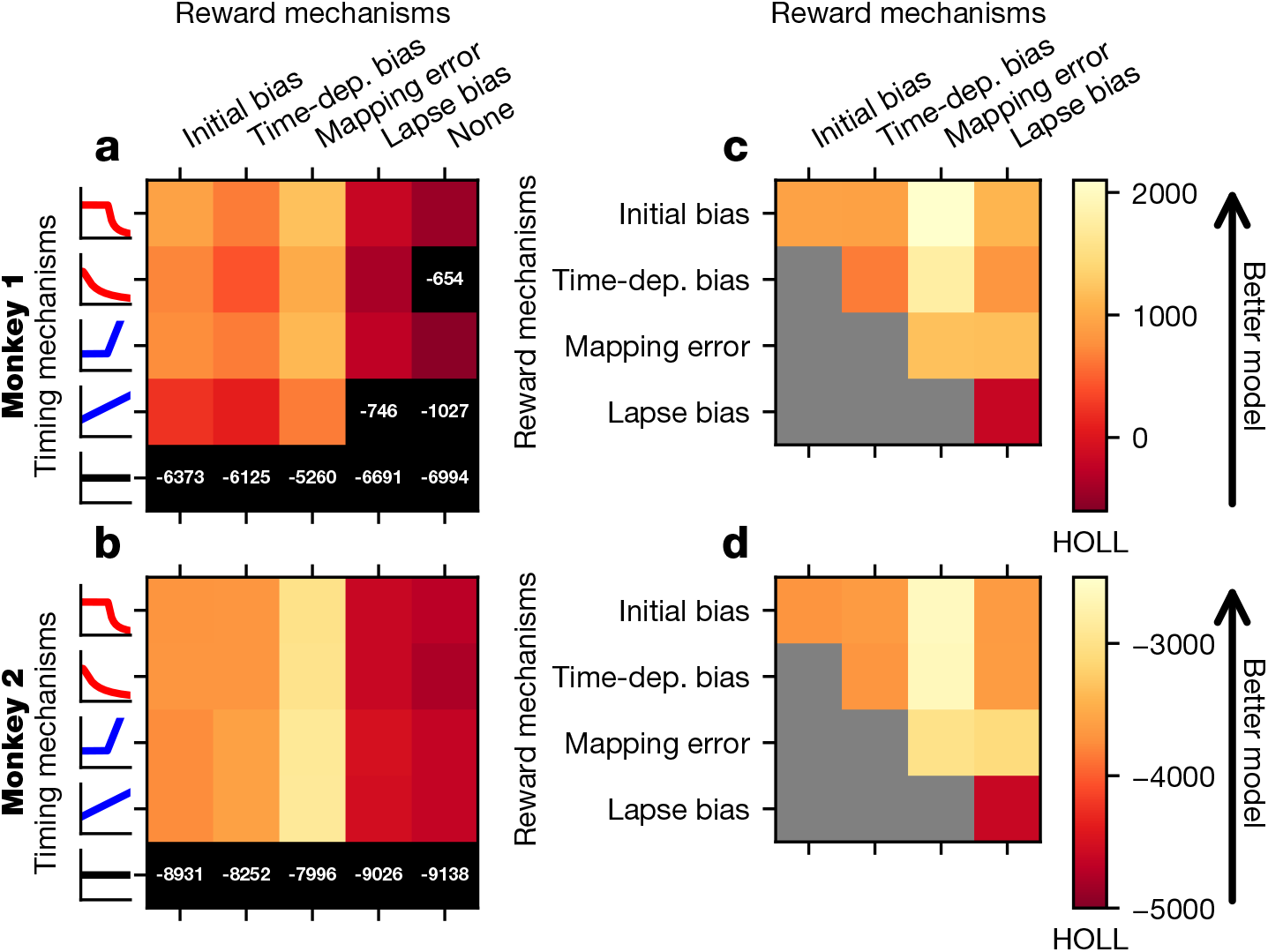
Fit of potential GDDM models. Various models were fit to the exploration data from monkey 1 (a,c) or monkey 2 (b,d) and subsequently evaluated with HOLL on the validation data. Color indicates the HOLL of the model, with higher values indicating a better fit. (a,b) Models were constructed by selecting one urgency signal (***Figure 3***b) and one reward mechanism (***Figure 3***a). Models with HOLL outside the range of the color bar are shown in black with their corresponding overlaid HOLL value. (c,d) Models were constructed using the delayed collapse urgency signal in conjunction with one or two reward mechanisms (***Figure 3***a).Models with only one reward mechanism appear on the diagonal. All models used leaky integration. **Figure 4–Figure supplement 1. Fit of GDDM models with exploration data.** Format similar to ***Figure 4***, except fit is evaluated using BIC on the exploration data instead of HOLL on the validation data.

Regarding reward bias mechanisms, we first compared GDDMs with only one of the four reward bias mechanisms, and found that the models with mapping error performed best. This suggests that asymmetric rewards affected the animal’s choice through an integration-independent mechanism (***Figure 4***a,b). Moreover, models with a time-dependent bias did not perform better than those with initial bias, suggesting that the reward bias mechanisms need not depend on time in order to explain the observed effects. We next asked whether these bias mechanisms could be combined to further improve model fit. In conjunction with a delayed collapsing bounds urgency signal, we tested all combinations of two reward bias mechanisms, and found that adding initial bias to the mapping error mechanism consistently improved the model fit for both monkeys more than any other mechanism such as time-dependent bias or lapse bias (***Figure 4***c,d). While initial bias and time-dependent bias both performed well individually and with mapping error, they did not improve the fit of the model when considered together (***Figure 4***c,d). These results suggest that both integration-dependent and integration-independent mechanisms are needed to explain animals’ reward bias.

We next examined the role of leak in evidence accumulation in the GDDM. For our best-fit model, the time constant was 140 ms for monkey 1 and 63 ms for monkey 2. We found that given urgency mechanisms, the estimated time scale of integration was consistently short (<150 ms) across all the tested models (***Figure 5***). Model comparison showed that adding a leak parameter to each model improved model fit for all the models except the simple DDM (***Figure 5***), for which the best-fitting leak time constant was often infinity. Notably, the models with non-delayed urgency and the models implementing time-dependent bias mechanism showed particularly large improvement with the addition of the leak parameter (***Figure 5***a,b). This result suggests that the leaky integration might improve the performance of these models by providing a mechanism to disregard early uncertain evidence, a property which can also be represented through a delayed urgency signal. However, the fact that the leak parameter also produced substantial improvement in delayed urgency models suggests that the short time scale of integration itself is an important feature.

**Figure 5.**
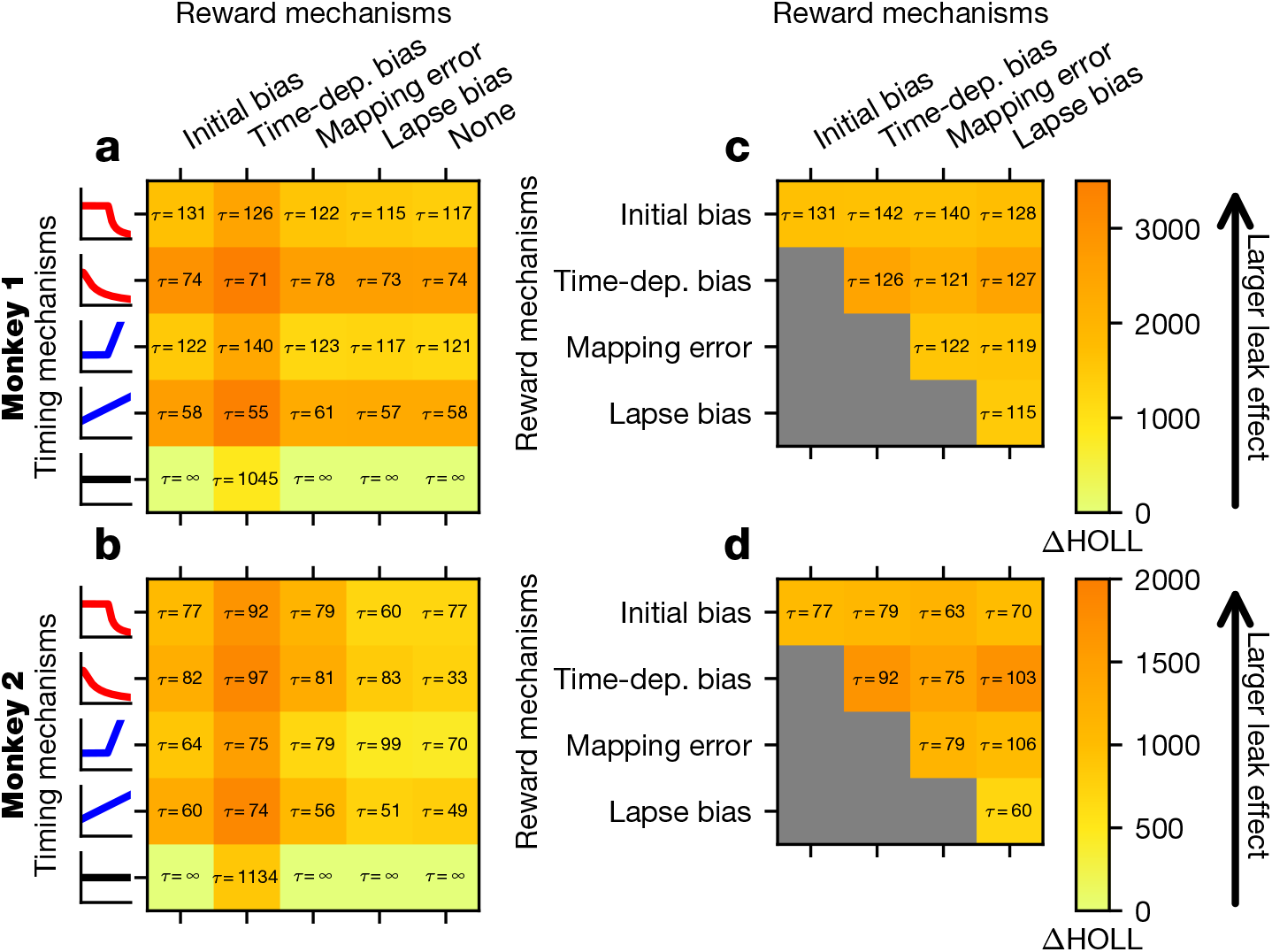
Leaky integration improves model fit non-uniformly. All models shown in ***Figure 4*** were subsequently fit to the exploration data with leaky integration disabled. The difference in HOLL on the validation data is shown, with larger ΔHOLL indicating a larger effect of leaky integration. As in ***Figure 4***, models involving one reward mechanism and one urgency signal (a,b) as well as models involving two timing mechanisms in conjunction with the delayed collapse urgency signal (c,d) were examined for monkey 1 (a,c) and monkey 2 (b,d). Overlaid are leaky integration time constants in units of milliseconds for each model. **Figure 5–Figure supplement 1. Leaky integration with exploration data.** Format similar to ***Figure 5***, except fit is evaluated using BIC on the exploration data instead of HOLL on the validation data.

The best-fit model captured behavior quite well. It fit the psychometric and chronometric functions and exhibited major features of choice and RT data that cannot be accounted for by simple DDM with integration-based reward mechanisms alone (***Figure 2***). In particular, the best-fit model dramatically improves the quantitative and qualitative fit to the RT distributions, compared to the simpler model (***Figure 2***). This model provided the best fit of all models considered here for Monkey 1, and nearly the best fit for Monkey 2, for which a slightly higher likelihood can be obtained through the use of the delayed gain function as an urgency signal (***Figure 4***).

As mentioned above, the RT distributions in our data during the low-coherence sample overlap with one another for all presample durations, and this could not be captured by the simple DDM with reward bias which uniformly integrates sensory evidence (***Figure 2–Figure Supplement 1***). By contrast, the best GDDM correctly predicts overlapping RT distributions as well as the RT-accuracy relationship for low-coherence samples (***Figure 2***). When evidence is weak, delayed urgency combined with a short time scale of integration implies the decision is most likely to be driven not by the total evidence, but rather by the later evidence close to the maximum presample duration used (0.8 s), i.e. when the uncertainty about stimulus onset is resolved.

Finally, the best GDDM fit the time course of errors frequently occurring after the onset of the high-coherence sample. This pattern of errors is inconsistent with one of the basic tenets of the DDM that stronger sensory evidence for one alternative makes the subject more likely to choose that alternative. To the contrary, our result shows that in the presence of asymmetric rewards, strong evidence for one alternative can paradoxically make the subject more likely to choose the other alternative. The apparently dynamic reward bias that increases with presample duration is mostly captured by the mapping error mechanism, indicating a failure of mapping the decision variable to motor output correctly.

### Manipulating temporal expectation changes strategy

Thus far, our findings suggest a set of mechanisms in the GDDM to quantitatively capture the behavioral choice and RT patterns. The two temporal features in this model, leaky integration and a delayed urgency signal, hint that the temporal structure of the task may drive their properties. By manipulating the task’s structure, we tested whether the urgency signal could be flexibly influenced by temporal context within a single experimental session.

We investigated this question using a variant of the task in the same monkeys (***Figure 6***). In this new task, the duration of temporal uncertainty was manipulated by varying the set of alternative presample durations between different blocks of trials within a session. In the “short-presample” blocks, there was a 0.5 s presample in majority of the trials (60%); in the remaining 40% of trials, the sample arrived either earlier (0.25 s) or later (0.75 s) with 20% probability each. Similarly, in the “long-presample” block, 60% of trials had a presample of duration 1.25 s, and 20% each had 0.75 s and 1.75 s (***Figure 6***). Accordingly, 20% of trials in each block had a 0.75 s presample, which corresponded to the longest presample duration in the short-presample block and the shortest presample duration in the long-presample block. No reward bias was imposed in this modified version of the task, so a correct response to either target elicited an identical reward.

**Figure 6.**
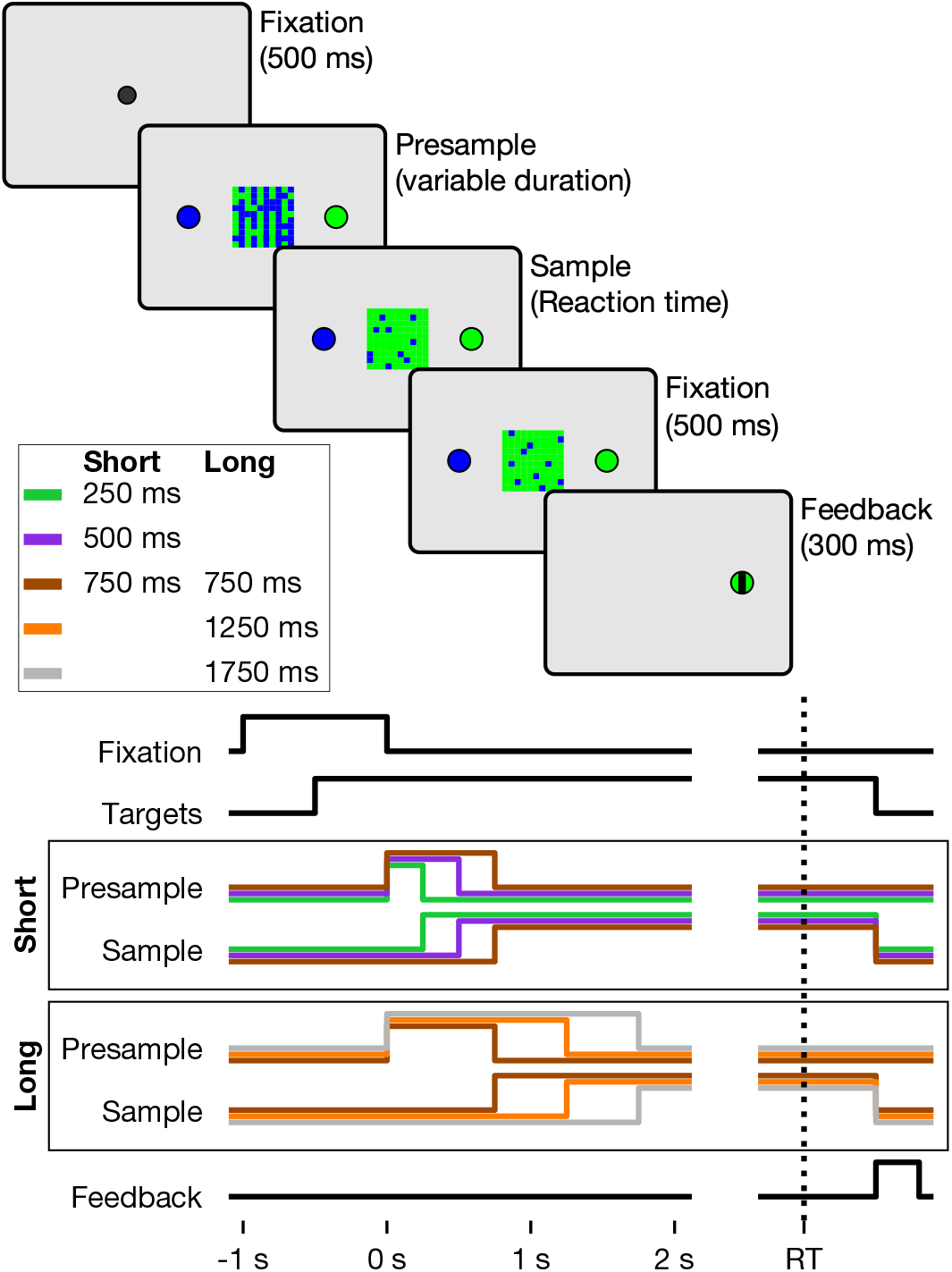
Color matching task with timing blocks. The temporal sequence of trial events in the color matching task with timing blocks is shown above. Below is a timeline indicating the presence of various task elements on the screen for either the short- or long-presample blocks, with presample and sample lines for each condition denoted by the line color.

This task was utilized to test whether temporal context can modify the urgency mechanisms of the fitted GDDM. Under a pure DDM framework, the RT distribution for the 0.75 s presample should be the same in both blocks. However, if temporal context influenced behavior, the RTs for the 0.75 s presample in the short-presample block would be on average shorter than those in the long-presample block.

We first performed a linear regression analysis in order to determine the effect of coherence, presample duration, and block on RT and accuracy (see Methods). As expected, we found higher coherences and shorter presample durations both decreased RT (*p* < .05, ***Equation 5***) and increased accuracy (*p* < .05, ***Equation 6***). More importantly, we also found a significant effect of the block regressor, in that trials presented during the long-presample blocks had significantly longer RT (*p* < .05, ***Equation 5***) and higher accuracy (*p* < .05, ***Equation 6***) compared to the short-presample blocks.

To examine more mechanistically how the animals altered their decision-making strategies, we fit the best GDDM chosen for the first task, including a leaky delayed collapsing bound, to the second task. Mechanisms for reward bias were removed given the lack of reward asymmetry in the present task. This model provided an excellent fit to the psychometric and chronometric functions as well as the probability distribution (***Figure 7***).

**Figure 7.**
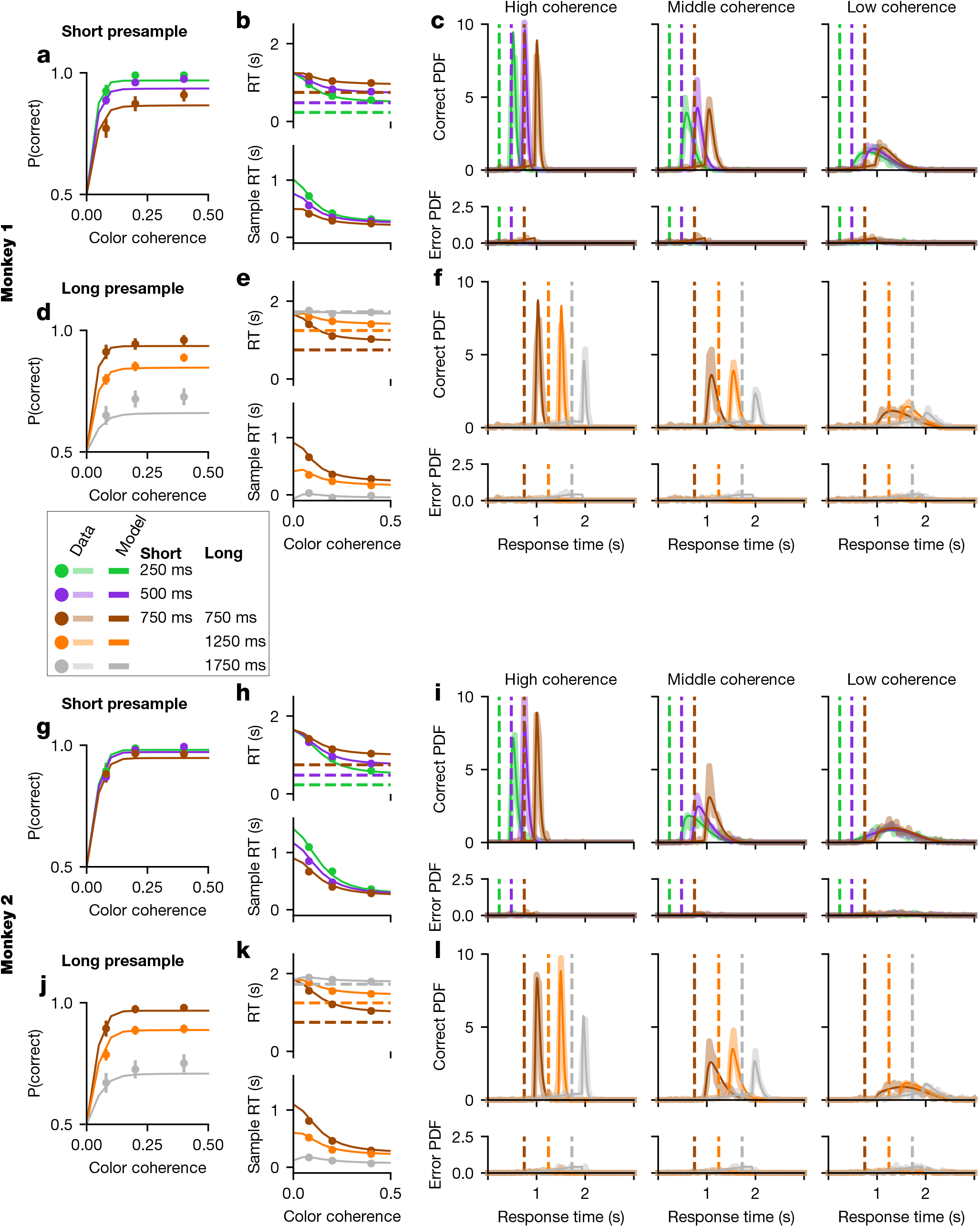
Model fit to timing dataset. Psychometric functions (a,d,g,j), chronometric functions (b,e,h,k), and RT histograms (c,f,i,l) are shown for short-presample (a,b,c,g,h,i) and long-presample (d,e,f,j,k,l) blocks for monkey 1 (a,b,c,d,e,f) and monkey 2 (g,h,i,j,k,l). Data are represented by dots in the behavioral functions and thick translucent lines in the probability distributions. “Sample RT” indicates the RT minus the presample duration. Overlaid as a thin solid line is a GDDM including delayed collapsing bounds and a leaky integrator with a separate “collapse delay” *t*_1_ parameter for short- and long-presample blocks. Error bars for 95% CI are hidden beneath the markers.

To investigate which model parameters could best explain the difference between blocks, we allowed individual parameters in this model to differ for the two blocks and assessed the log-likelihood in each case. For comparison, we also fit a 16-parameter model in which all of the parameters were allowed to vary between blocks, and an 8-parameter model which fixed all parameters to be the same for both blocks.

We examined the difference in HOLL when each parameter was fit separately for each block compared to shared between the blocks, as well as when all parameters or no parameters are shared. We found that the 16-parameter model fit the data better than models in which only a single parameter was allowed to vary between the short and long blocks. Nevertheless, changes in the urgency signal alone were sufficient to explain most of this improvement in model fit (***Figure 8***). For example, in ***Figure 7***, the model which allows the “collapse delay” *t*_1_ parameter to vary is shown. This demonstrates that the timing of the urgency signal mediates a critical aspect of the monkey’s change in strategy between different temporal contexts.

**Figure 8.**
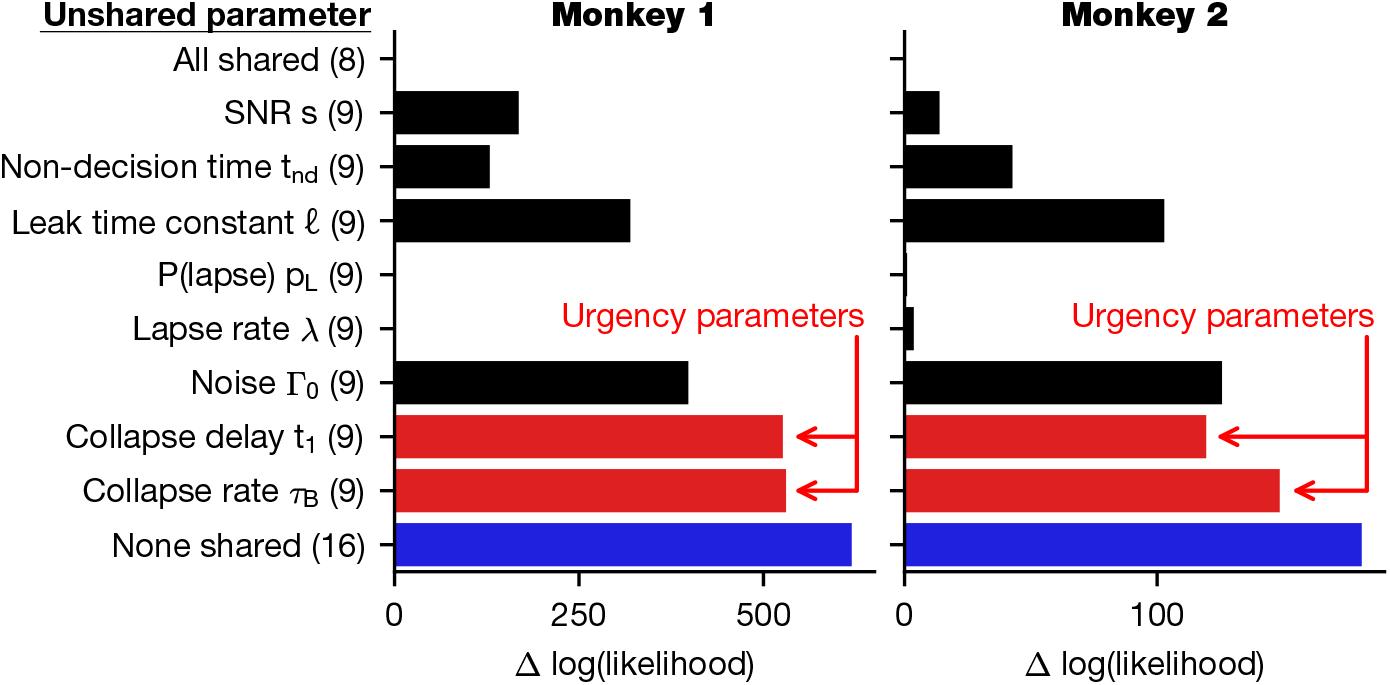
Urgency signal captures changes between presample blocks. An eight-parameter GDDM with a delayed collapse urgency signal and leaky integration was fit to data from the color matching task with presample blocks. The block type (either long- or short-presample) was incorporated into the model by allowing one parameter at a time to vary between blocks, forming a 9 parameter model, and then re-fitting all parameters of the new model to the data. For comparison, we also fit the 16 parameter model where we allowed all parameters to differ between blocks (blue). The improvement in log likelihood for each model is shown, as well as the total number of parameters of the model in parentheses. Monkey 1 is shown on the left, and monkey 2 on the right. Parameters related to the urgency signal are highlighted in red.

### Urgency timing can implement a speed-accuracy tradeoff

The above results showed that context-dependent changes in the timing of the urgency signal can account for the changes in behavior related to temporal uncertainty. In theory, the animal could increase accuracy by simply delaying the onset of the increase in urgency signal for the longest possible duration (1750 ms) on every trial, which would minimize the effect of integrating noise. However, this would come at the cost of long RT. Conversely, the animal could reduce RT by beginning to increase urgency immediately at the beginning of the trial, but at the cost of lower accuracy. Therefore, the onset of the urgency signal can mediate a speed-accuracy tradeoff, with longer onset delays favoring accuracy and shorter onset delays favoring speed.

To examine this quantitatively, we systematically varied the “collapse delay”*t*_1_ (***Figure 3***b) from 0 ms to 1000 ms, separately for long- and short-presample blocks, to observe how this parameter modulated the speed-accuracy tradeoff in the GDDM, with all other parameters shared (***Figure 9***). This analysis confirmed that changes to the urgency signal are able to control the speed-accuracy tradeoff. Furthermore, the speed-accuracy tradeoff is strikingly similar for both monkeys, despite the differences in estimated parameter values. This demonstrates that strategic modulation of speed-accuracy tradeoff can be accomplished using changes to only the urgency signal, which suggests that urgency may be critical for timing-related decision strategies.

**Figure 9.**
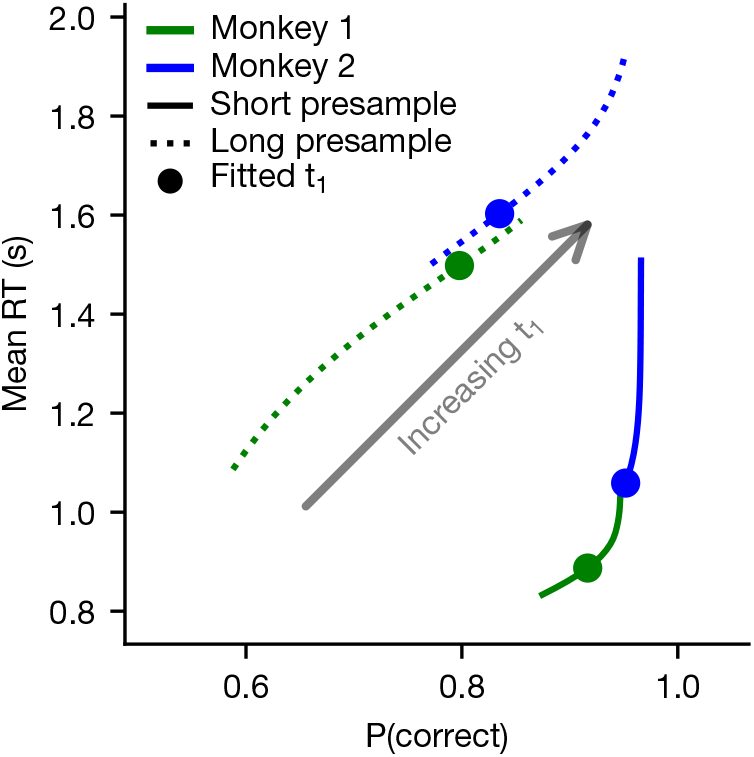
Urgency signal mediates a speed-accuracy tradeoff. A GDDM with a delayed collapse urgency signal and leaky integration was fit to each monkey’s RT data in the color matching task with presample blocks. Using these parameters, the “collapse delay *t*_1_” parameter was varied systematically across the linearly-spaced values from 0 to 1, representing the hypothetical case in which the subject could control this parameter while leaving the other parameters fixed. For each value, the mean RT and probability of a correct response were plotted for the long-presample (dotted lines) and short-presample (solid lines) blocks. The actual parameters are shown as filled dots.

## Discussion

In this study, we found that perceptual decision-making is driven by temporal changes in evidence quality via a dynamic urgency signal. We also showed that reward bias in animals’ choice can be accounted for by integration-dependent and integration-independent mechanisms.

### Temporal uncertainty and decision-making

An urgency to commit to a decision has long been hypothesized to modulate the speed and accuracy of perceptual decision-making (***Reddi and Carpenter, 2000***). In computational models, urgency has been implemented either by a collapsing bound or gain modulation of sensory evidence (***Ditterich, 2006a,b; Forstmann et al., 2008, 2010; Ratcliff and McKoon, 2008; Churchland et al., 2008; Cisek et al., 2009; Drugowitsch et al., 2012; Hanks et al., 2014; Murphy et al., 2016***). In models with collapsing bounds, an evidence-independent, time-varying signal is combined with the accumulators with fixed weight on evidence, while the gain function multiplicatively modulates the evidence with time-varying weight. However, the nature of time-varying mechanisms has been controversial (***Hawkins et al., 2015a,b; Voskuilen et al., 2016***). During the task used in our study, we varied the uncertainty about evidence onset over time to probe the time-varying mechanisms. Using an array of computational models, we showed that an urgency signal provides a flexible mechanism by which temporal information can be incorporated into the decision.

Recently, several studies have found, using dynamically changing sensory evidence, that subjects can adjust the gain or weight on the evidence across time during the trial, depending on the temporal statistics of evidence (***Cheadle et al., 2014; Levi et al., 2018***). In these previous studies, however, subjects were not allowed to freely choose the timing of their response, so it was not possible to determine how the subject’s strategy reflected a speed-accuracy trade-off. In the present study, we adopted an RT paradigm and explicitly showed that animals voluntarily adjusted the timing of their decision in order to reduce uncertainty about evidence through task-specific timevarying urgency. While we did not explore the potential optimality of this strategy, this presents an interesting opportunity for future study (***Drugowitsch et al., 2012, 2014***).

Time-varying stimuli were also used to test the time scale of evidence integration (***Bronfman et al., 2016***). Whereas the traditional DDM assumed a perfect integrator, recent studies have raised possibility that evidence integration may be leaky, as the accuracy of the decision does not necessarily increase with integration time (***Usher and McClelland, 2001; Zariwala et al., 2013; Tsetsos et al., 2015; Farashahi et al., 2018***). Moreover, when evidence dynamically changes during the course of a trial, models with leaky integration or instantaneous evidence fit the data well, as shown in the present study (***Cisek et al., 2009; Thura et al., 2012; Thura and Cisek, 2014; Bronfman et al., 2016***). In addition to leaky integration, bounded integration has also been proposed as a mechanism to account for the apparent failure to use the full stream of stimulus information (***Kiani et al., 2008***).

Our use of a dynamic urgency signal can be seen as a generalization of the idea that integration must begin at a single point in time (***Teichert et al., 2016***). Recently, integration onset was examined by ***Devine et al. (2019)***, who tested human subjects on a perceptual decision-making paradigm with multiple presample durations, a single sample coherence level, and no reward bias. Based on behavioral and EEG data, they proposed that under temporal uncertainty, integration begins approximately at the time of the earliest sample onset. However, their analyses did not fit a model of decision-making processes to test against alternative mechanisms. By contrast, we find strong evidence that our subjects used the temporal statistics of the task to modulate the evidence accumulation process. Applying our model comparison framework in other task paradigms can test the generality of these strategies.

A monotonically increasing gain function provides a straightforward mechanism to weight late evidence more than early evidence. The collapsing bounds mechanism does not by itself weight late evidence more than early evidence, although the interplay between time-varying urgency and leaky integration may provide a mechanism for temporal weighting of late evidence over early evidence. For example, in fixed-duration paradigms, a leaky integrator can effectively weight late evidence more than early evidence (***Levi et al., 2018; Lam et al., 2017***). We found that leaky integration improved fit in all models but played a larger role for simpler urgency signals than for the best-fitting delayed urgency signals, suggesting that leaky integration may capture a related component of behavior as the delay in urgency increase. While our results imply that both leaky integration and a delayed urgency signal are necessary, the ability of a time-varying urgency signal to reduce the relative benefit of adding leaky integration to the model demonstrates the flexibility of the urgency signal.

### Asymmetric rewards and decision-making

Asymmetry in reward or prior probability of the correct target can induce a bias to targets with a higher expected value (***Voss et al., 2004; Lauwereyns et al., 2002; Diederich and Busemeyer, 2006; Ratcliff and McKoon, 2008; Mulder et al., 2012; Feng et al., 2009; Rorie et al., 2010; Teichert and Ferrera, 2010***). Several studies have indicated that reward bias in perceptual decision-making could be well accounted for by an integration-dependent mechanism in which reward is modeled either a constant evidence (i.e. initial bias) or momentary evidence (i.e. time-dependent bias) (***Feng et al., 2009; Rorie et al., 2010; Fan et al., 2018; Gesiarz et al., 2019***). In our task, even a combination of these two reward mechanisms could not fit our data in the absence of an urgency signal or leaky integration. Likewise, recent studies have reported that under time pressure, reward bias might be produced by integration-independent mechanisms such as a quick guess that races with the process of evidence integration (***Simen et al., 2009; Noorbaloochi et al., 2015; Afacan-Seref et al., 2018***). However, this possibility was not consistent with our data either. This discrepancy might be due to the fact that previous studies have seldom tested the possible role of urgency in modulating the balance between evidence and bias when determining choice. Although reward bias increasing with time following the onset of an informative sensory stimulus could imply a time-varying bias mechanism, our results showed that such a time-varying bias is not necessary to explain our data. Overall, our results suggest that temporal and reward structures play an important role in perceptual decision-making. In addition, models within the GDDM framework can effectively describe the contribution of temporal and reward related factors during decision-making and play an important role in investigating the cognitive mechanisms of decision-making.

As investigators begin to explore more complicated behavioral tasks and computational models, it is important to use methods of model-fitting which are specialized for complex, high-dimensional models. In this study, we fit each model to the entire RT distributions of correct and error choices using maximum likelihood, and were able to provide qualitatively good fits to both the RT distribution and the psychometric and chronometric functions. By contrast, much previous work has fit models directly to the psychometric and chronometric functions, producing excellent fits to these reduced behavioral functions with a potentially poor fit to the RT distributions. Indeed, we found that evaluating and providing model fits to the entire probability distribution are highly informative for differentiating performance of different models and mechanisms. This methodology is imperative for simulating and fitting complex models. Our methodological innovations will therefore allow for increased complexity in experimental design to test a broad range of hypotheses about the mechanisms of decision-making.

## Methods

### Animal preparation

Two male rhesus macaque monkeys (Q and P, identified as monkey 1 and 2 respectively: body weight, 10.5–11.0 kg) were used. The animal was seated in a primate chair with its head fixed and eye position was monitored with a high-speed eye tracker (ET49, Thomas Recording, Giessen, Germany). All procedures used in this study were approved by the Institutional Animal Care and Use Committee at Yale University, and conformed to the Public Health Service Policy on Human Care and Use of Laboratory Animals and Guide for the Care and Use of Laboratory Animals.

### Behavioral tasks

Animals were trained to perform a color matching task, in which the onset time of the sample was systematically manipulated by varying the duration of non-informative stimulus (presample) that preceded the sample. Two versions of the task were used. In the color matching task with asymmetric reward (***Figure 1***), either large or small magnitude of reward was randomly assigned for identifying the correct color of the sample at a given trial. Presample duration was randomly selected from three discrete values at each trial. In the color matching task with timing blocks (***Figure 6***), two sets of presample intervals were used and presented in alternating blocks of trials (i.e. short- and long-presample blocks). In each block, presample duration was randomly selected from three discrete values. The magnitude of reward was the same for all correct choices. Animals were first trained for the task with timing blocks and then for the task with asymmetric reward.

#### Color matching task with asymmetric reward

Each trial began when the animal fixated a central square (white, 0.6°×0.6°in visual angle) (***Figure 1***). After the 0.3 s fixation period, cues indicating magnitudes of reward available from two alternative choices were presented at the potential locations of saccade targets. Two sets of reward cues were used: in addition to those shown in ***Figure 1***, for some sessions, a circle was used for the large-reward target and a diamond for the small-reward target. Two nested squares in thin gray lines (or a circle) and a single square in thicker white lines (or a 45°-angled square) indicated larger and smaller reward, respectively. The ratio of reward magnitude was 2:3 and 1:4 for monkey 1 and 2, respectively. Following the reward-cue period (0.4 s), a green disk was presented inside one of the two reward cues and a blue disk appeared within the other, serving as targets for choice saccade. After 0.4 s of target period, a checkered square with equal numbers of green and blue pixels (20 ×20; 1.63°×1.63°in visual angle), or presample, was presented replacing the central fixation target. In addition, green and blue pixels were dynamically redistributed to random positions at a rate of 20 Hz. After a variable interval (0, 0.4, 0.8 s), the presample was replaced by the sample, which was identical to the presample except that it contained green and blue pixels in different proportions. Presample duration was randomly sampled from the three intervals with equal probability. The animal was allowed to shift its gaze towards one of the peripheral target at any time after the onset of the sample and was rewarded only when it correctly chose the target with the same color as the majority of the pixels in the sample. Choices made during presample were punished by an immediate end of the trial followed by a 2 s timeout. Three levels of color coherence were used for the sample (53%, 60%, 70% for monkey 1; 52%, 57%, 63% of pixels for monkey 2), each with equal probability. In about 5% of the trials, the presample never changed to a sample, meaning the coherence of each color was 50% and there was no correct choice. On these trials, a reward was delivered with 50% probability regardless of the monkey’s choice. After 0.3 s of fixation on the chosen target, feedback was provided for 0.3 s with a horizontal or vertical bar superimposed on the chosen target indicating erroneous or correct choice, respectively. Reward was delivered for correct choices. Color coherence, location of saccade target with correct color, presample duration, and location of large-reward cue were pseudo-randomly selected for each trial. Monkey 1 performed 30868 trials across 50 sessions, and monkey 2 performed 23695 trials across 29 sessions.

#### Color matching task with timing blocks

The sequence of trial events was the same as in the color matching task with asymmetry (***Figure 6***), except that fixation and saccade targets were presented simultaneously without the reward cue, and the durations of the presample varied between blocks. Presample duration for the majority of the trials (60%) in the short- and long-presample block was 0.5 s and 1.25 s, respectively. In the remaining trials, the presample was terminated with equal probability (20%) either earlier (0.25 s and 0.75 s for short- and long-presample blocks, respectively) or later (0.75 s and 1.75 s for short- and long-presample blocks, respectively) than the most frequent presample durations. Choices made during the presample were penalized by subsequent 5 s timeout. Three levels of color coherence were used for the sample (54%, 60% and 70% for both monkey 1 and 2), each with equal probability. Short- and long-presample blocks alternated within a given session and the order of the two types of block was randomized across sessions. Monkey 1 performed 12,773 trials across 10 sessions, and monkey 2 performed 10,777 trials across 10 sessions.

### Fitting method

As we fit models with many parameters, we must be careful about the methods used for parameter estimation. One common approach in the field is to fit the parameters of the DDM based only on the probability of a correct response and the mean RT for correct trials. This uses only a very small subset of the data, using only two summary statistics, and is thus insufficient for fitting complex models. A second approach which is commonly used when fitting complex models is to simulate individual trajectories of the decision variable, and then perform tests on the quadrants of the simulated values to determine whether there is a match. While this approach utilizes more data than the previously discussed method, it still uses only summary statistics of the data. More critically, this simulation process is very slow, and hence does not permit efficiently fitting the model to data. These two approaches are principally used to overcome limitations in simulating and fitting these models, namely, that a full RT distribution is not always available. Our approach simulates the stochastic differential equations by numerically solving the Fokker-Planck equations so that we may use likelihood-based methods on the entire RT distribution for estimating parameters. This allows all data to be utilized through a robust statistical framework.

RT distributions simulated directly from the Fokker-Planck equation often use the forward Euler method, an easy-to-implement method for solving stochastic differential equations. The forward Euler method mandates the use of very small time steps and a coarse decision variable discretization to maintain numerical stability. In practice, this means that simulations which achieve a reasonable margin of error are prohibitively time-consuming for fitting parameters.

To circumvent this, we instead solve the Fokker-Planck equation using the Crank-Nicolson method (for the fixed-bound conditions) or the backward Euler method (for the collapsing bound conditions). These methods do not require small time steps in order to achieve low margin of error. As a result, we are able to fit parameters using the results of these numerical solutions. Simulations were performed using the PyDDM package^1^ using a timestep of 5 ms and decision variable discretization of 0.005. Correctness of the implementation was verified using specialized techniques (***Shinn, 2019***).

Fitting is performed by maximizing the log-likelihood function. For low-dimensional models, parameters may be optimized using a variety of local search functions which tend to give similar results. However, for high-dimensional models, these methods are often unable to minimize the function, instead finding a local minimum or failing to converge. Because we were simultaneously fitting up to 16 parameters, using an appropriate optimization routine was critical. We used differential evolution, a heuristic search method as implemented in Scipy (***Jones et al., 2001***). Differential evolution is a global search method, meaning it is intended to avoid local minima in the search space, and has been shown to exhibit consistent results in practice (***Storn and Price, 1997***).This gave consistent parameter estimates and similar log-likelihoods for our GDDMs.

### Cross-validation procedure

Once obtaining a fit, there are many different processes which may be used for evaluating the fit of a model to protect against overfitting. There are two types of overfitting which are important to consider for complex models such as ours with many parameters. First, overfitting may occur because the model mechanism is complicated and thus the parameters of the model may fit to noise within the data. Second, overfitting may be the result of the modeler testing many potential model mechanisms and choosing the best mechanism. Simple models with few parameters may only be concerned with the first of these, but models with complicated mechanisms must guard against both types of overfitting.

Many papers simply evaluate the best fit and report these values. This does not protect against either overfitting the parameters or overfitting the model mechanism. A more sophisticated approach is to utilize a measure such as AIC or BIC, which takes the model complexity into account when comparing two competing models. However, these metrics judge model complexity exclusively by the number of parameters, and this has been shown to be a poor metric for evaluating model complexity (***Piantadosi, 2018; Myung et al., 2000***). Additionally, this only helps with over-fitting the parameter values and does not prevent overfitting of the model mechanism. A better approach is to utilize n-fold cross validation, which will properly penalize complex models over simple models by ensuring that the parameters are not overfit. However, this still leaves open the possibility for overfitting the model mechanism. For complex models, the best way to prevent overfitting is to use held-out data which is not analyzed or otherwise examined in any way until the final model has been constructed and the conclusions of the model have been determined.

Therefore, in order to ensure robustness and protect against both types of overfitting, we split the dataset into two halves, an “exploration” dataset and a “validation” dataset. All analyses and model fitting were performed only on the exploration data using Bayesian information criterion (BIC) for comparing models. The validation trials were not fit to the model or otherwise examined until all analyses for the present manuscript were complete. After unmasking the validation trials, no additional analyses were performed, no additional results were added to the manuscript, and no existing results were removed. These analyses are shown as supplementary figures (***Figure 2–Figure Supplement 2, Figure 4–Figure Supplement 1, and Figure 5–Figure Supplement 1***).For a more robust analysis, we evaluate models on the validation dataset using the held-out log-likelihood (HOLL), i.e. the likelihood under the held-out (validation) data when parameters were fit using the exploration dataset (***Figure 2, Figure 4, Figure 5***). All quantitative and qualitative results found using BIC were unchanged when using HOLL with the validation dataset.

We did not perform this validation procedure on the task with timing blocks, as sample size was limited. Nevertheless, all models were developed on data from the task with asymmetric reward before being applied to data from the task with timing blocks.

### Behavioral analyses

In analyzing animals’ behavior, we measured response time (RT) relative to the presample onset. However, we occasionally refer to the animal’s response time relative to the onset of the sample (“Sample RT”).

A linear model was fit to examine the dependence of RT on task parameters:

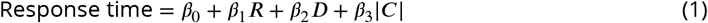

Here, *R* is reward magnitude, where 1 indicates that the large-reward target was the correct response and 0 that the small-reward target was correct; *D* is presample duration in units of seconds; and |*C*| is unsigned coherence, where 1 corresponds to a solid color patch and 0 to an equal ratio of each color. We define RT relative to the onset of the presample, rather than the sample, given that the transition from presample to sample was not explicitly cued. Coherence takes values greater than or equal to 0. All notation is described in ***Table 1***.

**Table 1.**
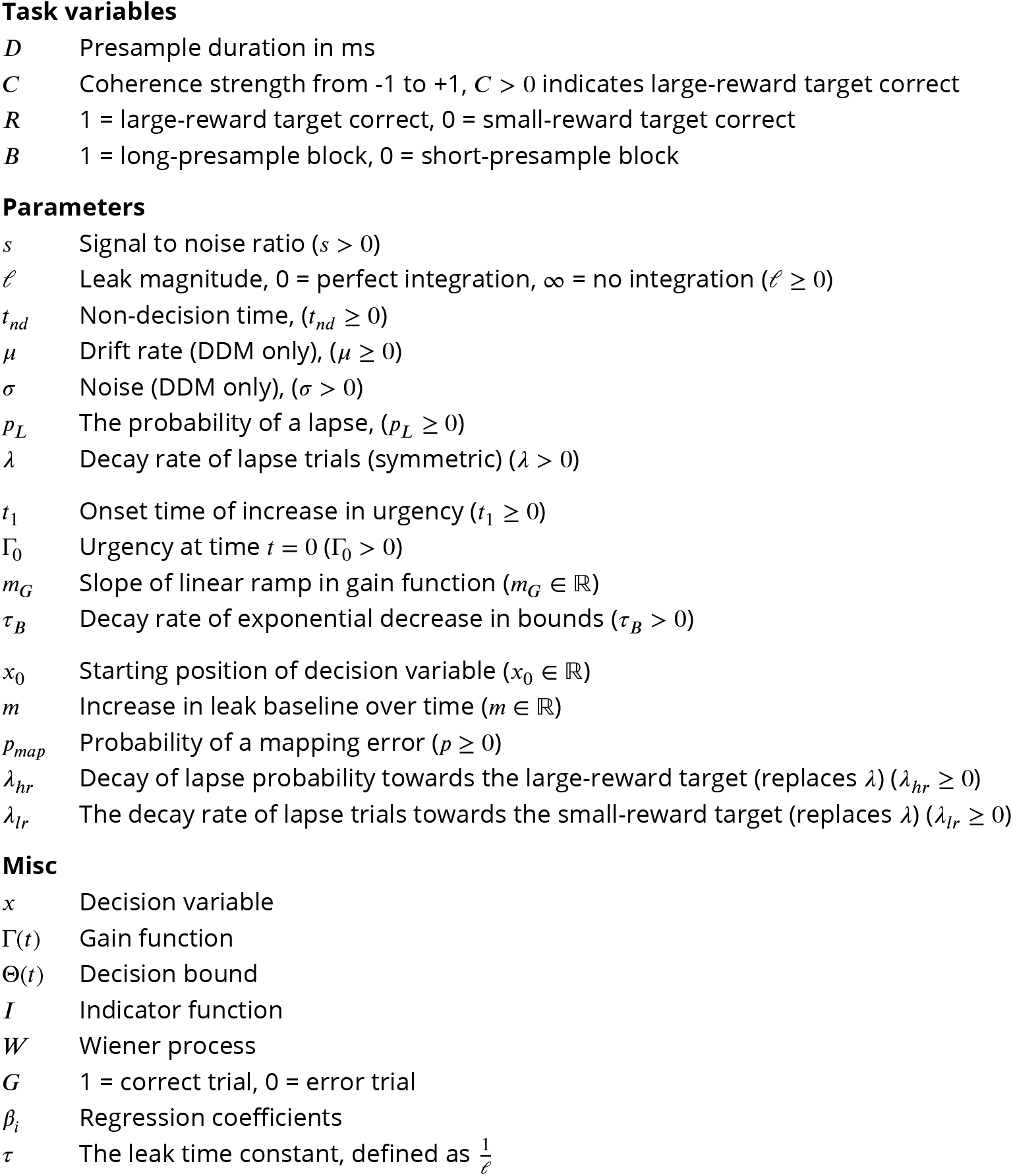
Summary of notation. All mathematical notation used in equations and models are listed below.

|  |  |
| --- | --- |
| $D$ | Presample duration in ms |
| $C$ | Coherence strength from -1 to +1, $C > 0$ indicates large-reward target correct |
| $R$ | 1 = large-reward target correct, 0 = small-reward target correct |
| $B$ | 1 = long-presample block, 0 = short-presample block |
| **Parameters** |  |
| $s$ | Signal to noise ratio ($s > 0$) |
| $\ell$ | Leak magnitude, 0 = perfect integration, ∞ = no integration ($\ell \geq 0$) |
| $t_{nd}$ | Non-decision time, ($t_{nd} \geq 0$) |
| $\mu$ | Drift rate (DDM only), ($\mu \geq 0$) |
| $\sigma$ | Noise (DDM only), ($\sigma > 0$) |
| $p_L$ | The probability of a lapse, ($p_L \geq 0$) |
| $\lambda$ | Decay rate of lapse trials (symmetric) ($\lambda > 0$) |
| $t_1$ | Onset time of increase in urgency ($t_1 \geq 0$) |
| $\Gamma_0$ | Urgency at time $t = 0$ ($\Gamma_0 > 0$) |
| $m_G$ | Slope of linear ramp in gain function ($m_G \in \mathbb{R}$) |
| $\tau_B$ | Decay rate of exponential decrease in bounds ($\tau_B > 0$) |
| $x_0$ | Starting position of decision variable ($x_0 \in \mathbb{R}$) |
| $m$ | Increase in leak baseline over time ($m \in \mathbb{R}$) |
| $p_{map}$ | Probability of a mapping error ($p \geq 0$) |
| $\lambda_{hr}$ | Decay of lapse probability towards the large-reward target (replaces $\lambda$) ($\lambda_{hr} \geq 0$) |
| $\lambda_{lr}$ | The decay rate of lapse trials towards the small-reward target (replaces $\lambda$) ($\lambda_{lr} \geq 0$) |
| **Misc** |  |
| $x$ | Decision variable |
| $\Gamma(t)$ | Gain function |
| $\Theta(t)$ | Decision bound |
| $I$ | Indicator function |
| $W$ | Wiener process |
| $G$ | 1 = correct trial, 0 = error trial |
| $\beta_i$ | Regression coefficients |
| $\tau$ | The leak time constant, defined as $\frac{1}{\ell}$ |

**Table 2.** Model parameters. Parameters for each displayed model. Rew: Color matching task with asymmetric reward, Time: Color matching task with timing blocks.

| Task: | Rew | Rew | Time | Time | Time | Time | Rew | Rew |
| --- | --- | --- | --- | --- | --- | --- | --- | --- |
| Model: | GDDM | GDDM | GDDM | GDDM | GDDM | GDDM | DDM | DDM |
| Monkey: | 1 | 2 | 1 | 1 | 2 | 2 | 1 | 2 |
| Block: | — | — | Short | Long | Short | Long | — | — |
| $s$ | 9.32 | 6.52 | 11.25 | 11.25 | 12.11 | 12.11 | | |
| $\Gamma_0$ | 1.03 | 1.16 | 1.62 | 1.62 | 1.40 | 1.40 | | |
| $t_{nd}$ | 0.22 | 0.21 | 0.20 | 0.20 | 0.22 | 0.22 | 0.27 | 0.21 |
| $\ell$ | 7.14 | 15.82 | 22.67 | 22.67 | 9.81 | 9.81 | | |
| $\tau_B$ | 1.19 | 0.70 | 0.35 | 0.35 | 0.47 | 0.47 | | |
| $t_1$ | 0.36 | 0.09 | 0.00 | 0.21 | 0.20 | 0.73 | | |
| $x_0$ | 0.09 | 0.05 | | | | | -0.16 | -0.04 |
| $p_{map}$ | 0.12 | 0.21 | | | | | | |
| $p_L$ | 0.04 | 0.03 | 0.03 | 0.03 | 0.04 | 0.04 | 0.03 | 0.10 |
| $\lambda$ | 1.02 | 0.12 | 0.47 | 0.47 | 1.47 | 1.47 | 0.48 | 0.95 |
| $\mu$ | | | | | | | 2.90 | 6.27 |
| $\sigma$ | | | | | | | 0.77 | 0.76 |
| $m_e$ | | | | | | | 0.60 | 0.43 |

To examine second-order interaction terms, a second model was fit:

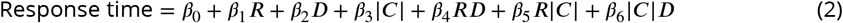

Similarly, a binomial generalized linear model was fit to understand how the task parameters influenced the monkey’s decision to choose the correct target:

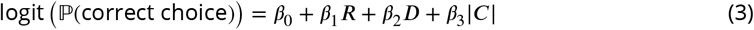

where variables are as defined above. Likewise, a second model was fit which contained second-order interaction terms:

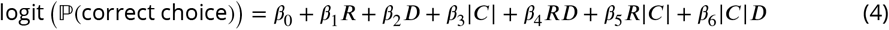

Linear model results were similar in both the exploration and validation datasets. Trials with zero coherence were excluded.

Likewise, RT was examined for the color match task with timing blocks using the linear model:

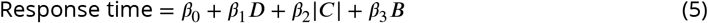

where *B* is the block, coded as 0 for the short-presample block and 1 for long-presample, and other terms are as specified above. Accuracy was likewise analyzed with the binomial generalized linear model:

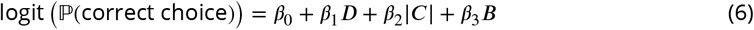

### DDM with reward mechanisms

The DDM is governed by a decision variable *x*. The evolution of *x* in our reward-biased DDM model, similar to that of ***Fan et al. (2018)*** and ***Gesiarz et al. (2019)***, is described by:

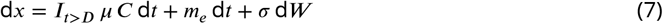

where the following parameters were fit to data:

- *μ* - signal strength
- *σ* - noise level
- *m_e_* - momentary sensory evidence bias

Additional terms in the model are:

- *W* - A Wiener process with standard deviation 1
- *C* - The color coherence
- *I*_*t*>*D*_ - An indicator function for the presample on the current trial. *I*_*t*>*D*_ = 0 if we are in the presample, and *I*_*t*>*D*_ = 1 otherwise.

We specify that *x* has initial condition given by parameter *x* = *x*_0_ at *t* = 0 seconds. We use the convention that positive values of *x* indicate choices towards the large-reward target, and thus positive coherence indicates motion towards the large-reward target and negative coherence away from it. When 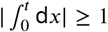, a decision to choose left or right is made based on the sign of *x*. Since the parameter *σ* is fit to data, integration bounds are fixed at 1 to prevent over-parameterization. The RTs generated by this model were shifted post-simulation by the non-decision time *t_nd_*.

Since we fit based on maximum likelihood, rare trials in which the choice appears unrelated to the stimulus (lapses) may have a large impact on the fitted parameters. Thus, for the purposes of model fitting, we fit a mixture model between the drift-diffusion process and an exponentially-distributed lapse rate with rate parameter *λ*. This is given by:

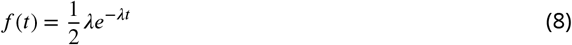

where there is a *p_L_* probability of any given trial being a lapse trial. While we fit both of these parameters to data, results were unchanged when we fixed these parameters at constant values (data not shown).

Thus, overall, this model contained seven parameters: *μ*, *σ*, *x*_0_, *m_e_*, *t_nd_*, *p_L_*, and *λ*.

### Generalized DDM

Like the DDM, the GDDM is governed by the evolution of a decision variable *x*. The instantaneous value of *x* is described by:

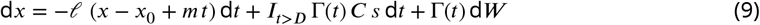

where the parameters which could potentially be fit by simulations are:

- *ℓ* - The leak parameter, constrained to *ℓ* ≥ 0. Its inverse is the leak time constant.
- *x*_0_ - The initial position of the integrator, and the location to which the leaky integrator decays. This was constrained to be *x*_0_ ≥ 0. By default, it was fixed to *x*_0_ = 0.
- *m* - The change in the leaky integrator baseline over time. By default this was fixed at *m* = 0.
- Γ(*t*) - The sensory gain, a function of time. By default, Γ(*t*) = Γ_0_.
- *s* - The signal-to-noise ratio, constrained to *s* ≥ 0.
- Θ(*t*) - The decision bound. When 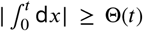, a decision to choose left or right is made based on the sign of *x*. Since noise is allowed to vary, integration bounds are subject to the constraint Θ(0) = 1 to prevent overfitting. In the absence of collapsing bounds, this fixes the bounds to be equal to ±1, the default.

Additional terms in this equation are defined as:

- *W* - White noise, i.e. a Wiener process.
- *C* - Coherence. We use the convention that positive values of *x* indicate choices towards the large-reward target, and thus positive coherence indicates motion towards the large-reward target and negative coherence away from it.
- *I*_*t*>*D*_ - An indicator function for the presample on the current trial. *I*_*t*>*D*_ = 0 if we are in the presample, and *I*_*t*>*D*_ = 1 otherwise.

As above, we fit a mixture model between the drift-diffusion process and an exponentially-distributed lapse rate with rate *λ*, with a *p_L_* probability of any given trial being a lapse trial. This is given by:

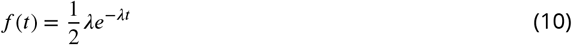

Overall, in the simplest possible case where all parameters are set to their defaults (thereby indicating that no reward or timing effects are included in the model), the model includes six parameters: *λ*, *p_L_*, Γ_0_, *ℓ*, *s*, *t_nd_*. Reward and timing mechanisms added to the model increased the number of parameters which were fit.

#### Reward mechanisms

Within the GDDM framework, we designed reward mechanisms for the model according to our hypothesized cognitive mechanisms, shown in ***Figure 3***.

#### Initial bias

The total evidence necessary to reach a decision is lower for the large-reward target than for the small-reward target. This is equivalent to changing the starting position of integration. Because our model deals with leaky integration, we furthermore impose that the leak decays to this position instead of to zero. This adds one parameter to the model: the magnitude of the baseline shift *x*_0_.

#### Time-dependent bias

The total evidence necessary to reach a decision is initially the same for the large- and small-reward targets, but a bias towards the large-reward target over the small-reward target develops linearly over time. Without leak, this is equivalent to a constant bias on the drift rate, i.e. a continuous integration of a constant. Because our model deals with leaky integration, such a bias would instead change the point to which the leaky integrator decays. Thus, we model this bias by temporally modulating the position to which the leaky integrator decays. This adds one parameter to the model: the slope of the increase in large-reward bias *m*.

#### Mapping error

After integrating to the bound, with some probability, small-reward choices are mapped to a response toward the large-reward target. Equivalently, some percentage of small-reward choices are assigned to be large-reward instead. This mechanism was first described in discrete evidence paradigms (***Erlich et al., 2015; Hanks et al., 2015***). This mechanism adds one parameter to the model: the probability of making a mapping error on any given trial *p_map_*.

Mapping error was implemented as post-simulation RT histogram modifications. For a simulated probability density function *f*(*x*|*R* = *r, C* = *c, G* = *g*, *D* = *p*), where *R* is large vs. small reward, *C* is coherence, *G* is correct or error trial, and *D* is the presample duration, we compute the final density *f*′(*x*) using *f*(*x*). For the mapping error, this was calculated as:

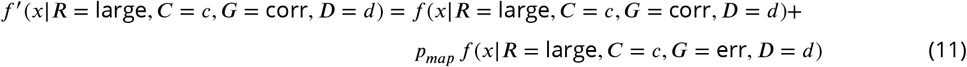

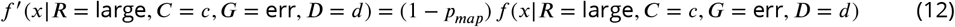

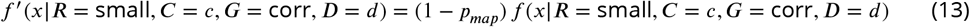

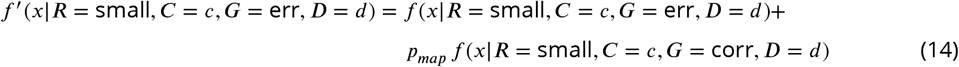

#### Lapse rate bias

The lapse rate, assumed to be exponentially distributed, is allowed to be higher in the direction of the large-reward target compared to the small-reward target. This is given for large- and small reward targets respectively by:

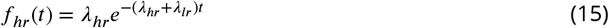

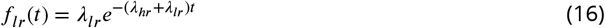

This mechanism has three parameters: the lapse rates for both large- and small-reward targets *λ_hr_* and *λ_lr_*, and the probability of any given trial being a lapsed trial *p_L_*. Models which include this mechanism do not include unbiased lapse rate, as it is redundant. Since the unbiased lapse rate uses two parameters, this mechanism adds one net parameter to the model.

#### Timing mechanisms

Similarly, we designed timing mechanisms to capture the ideas of urgency signals. Two types of urgency signal have been previously described in the literature: a “gain function” which scales evidence and noise uniformly throughout the course of the trial, and “collapsing bounds” which cause the decision bounds to become less stringent as the trial progresses.

#### Collapsing bounds

Collapsing bounds can be used to implement an urgency signal of various forms. While much debate has focused on the correct form of the bounds (***Hawkins et al., 2015a; Malhotra et al., 2017***), we use an exponentially collapsing bound for simplicity. This mechanism adds one parameter to the model: the time constant of the exponential collapse *τ_B_*. It is implemented as:

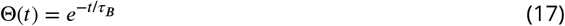

Note that this form satisfies the imposed boundary condition Θ(0) = 1.

#### Delayed collapsing bounds

This implements an exponentially decreasing bound as an urgency signal similarly to collapsing bounds, except the bounds do not begin to collapse until late in the trial. This mechanism adds two parameters to the model: the time before the bounds begin to collapse *t*_1_, and the rate of collapse *τ_B_*. This is implemented as:

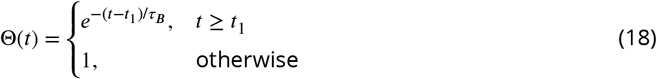

These also satisfies the boundary condition Θ(0) = 1 for all *t*_1_ ≥ 0.

#### Linear gain

A gain function can also be used to implement an urgency signal. It scales both the evidence and the noise simultaneously in order to preserve signal-to-noise ratio. This mechanism adds one parameter to the model: the rate at which gain increases *m_G_*. It is implemented as:

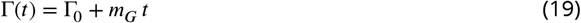

#### Delayed gain

The delayed gain function implements an urgency signal similarly to linear gain but it does not begin to increase the grain until part way into the trial. This mechanism adds two parameters to the model: the time at which the value begins to ramp *t*_1_, and the slope of the ramp *m_G_*. It is implemented as:

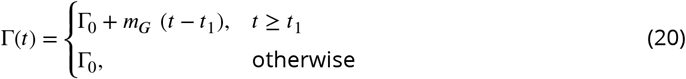

#### Simulations without leak

Simulations without leak were performed by setting *ℓ* = 0.

## Acknowledgments

We thank Norman H. Lam for assistance with model fitting.

---

**Figure 2–Figure supplement 1.**
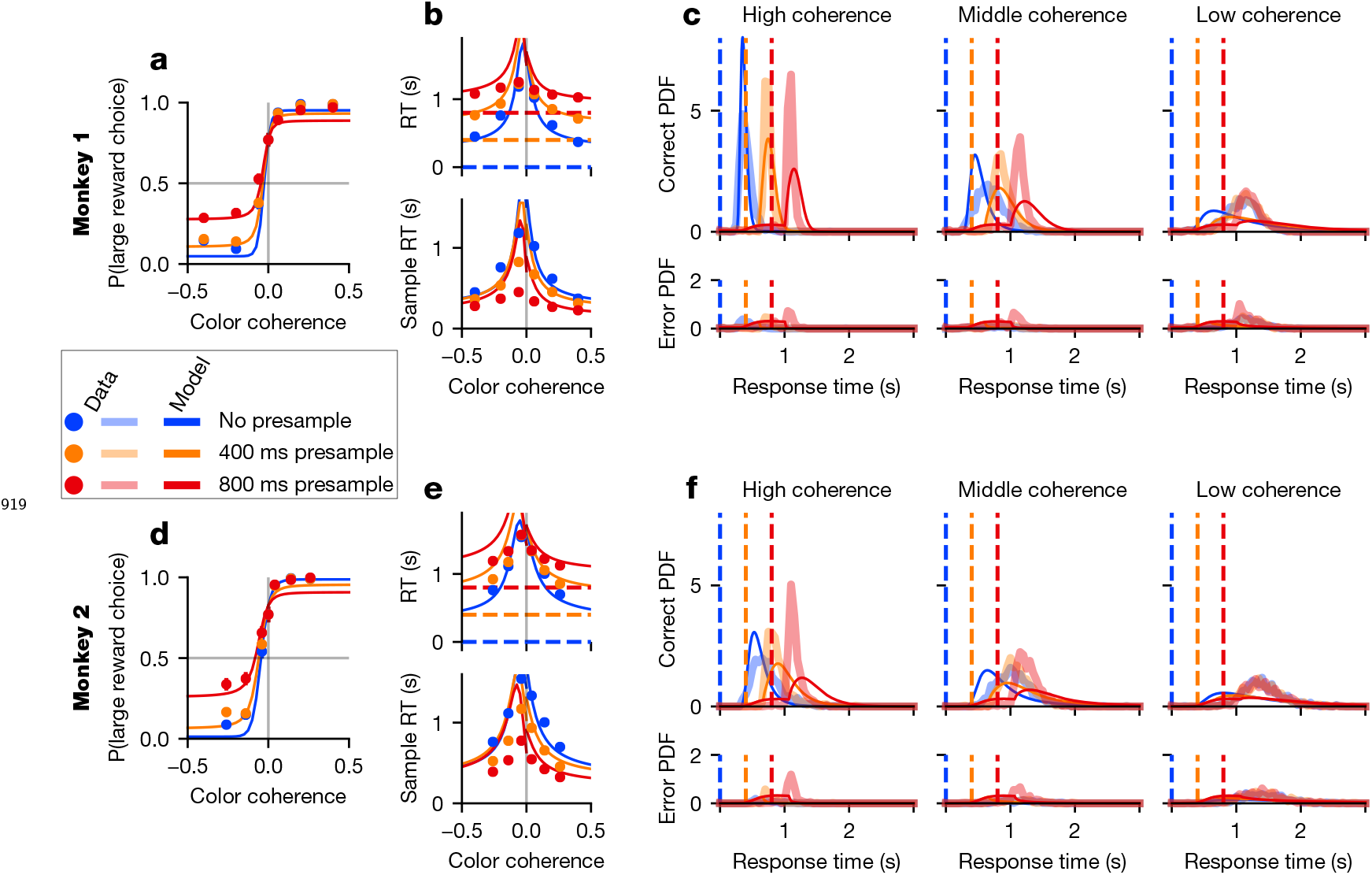
DDM with reward bias does not capture monkey behavior. Format similar to **Figure 2**, but with the model described in Drift-diffusion model with reward mechanisms.

**Figure 2–Figure supplement 2.**
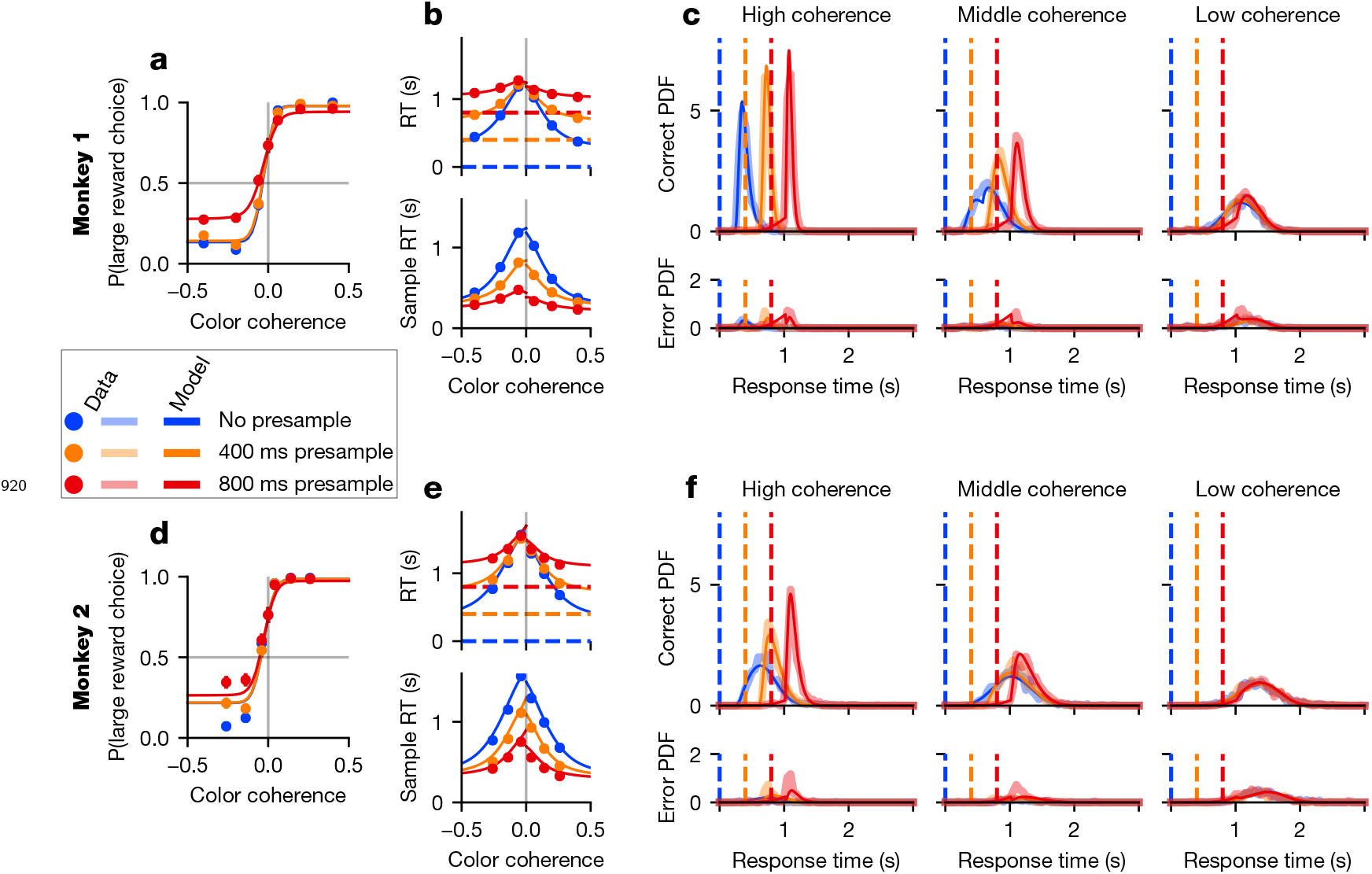
Animal and model performance in exploration dataset. Format similar to **Figure 2**, but showing the exploration data using model parameters fit to it.

**Figure 4–Figure supplement 1.**
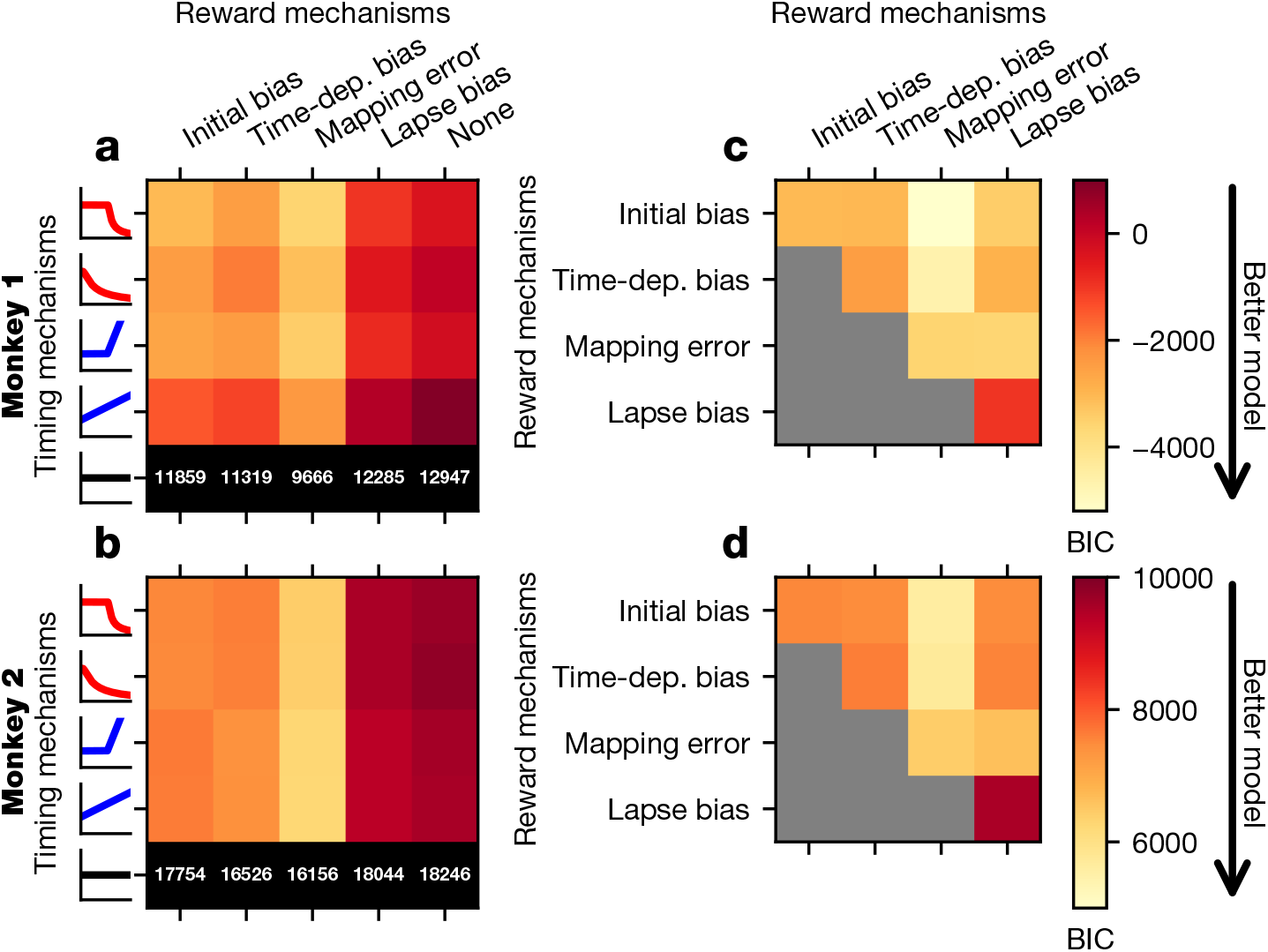
Fit of GDDM models with exploration data. Format similar to **Figure 4**, except fit is evaluated using BIC on the exploration data instead of HOLL on the validation data.

**Figure 5–Figure supplement 1.**
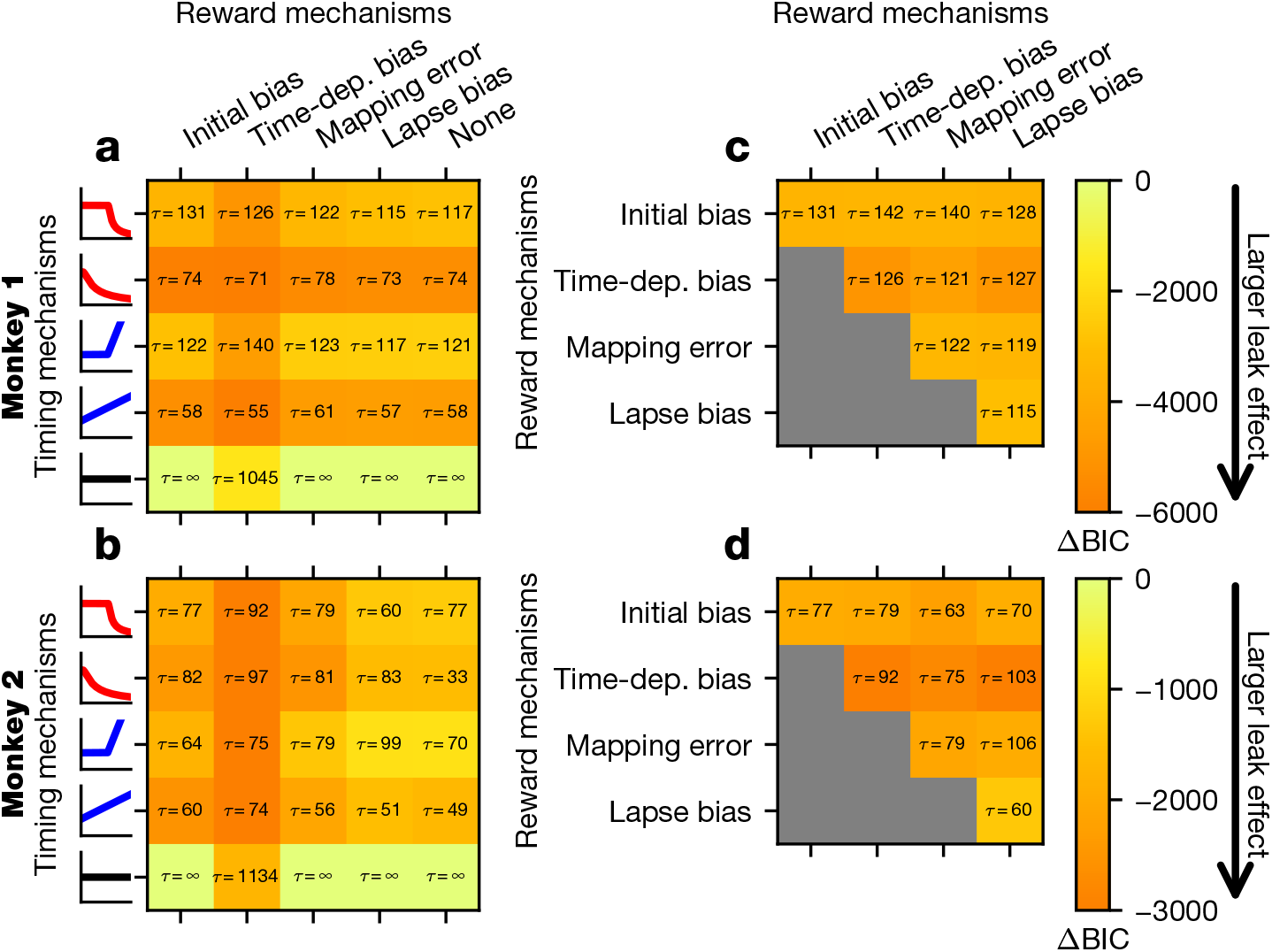
Leaky integration with exploration data. Format similar to **Figure 5**, except fit is evaluated using BIC on the exploration data instead of HOLL on the validation data.

## Footnotes

1 https://github.com/mwshinn/PyDDM

